# Wild cereal grain consumption among Early Holocene foragers of the Balkans predates the arrival of agriculture

**DOI:** 10.1101/2021.10.14.464411

**Authors:** E. Cristiani, A. Radini, A. Zupancich, A. Gismondi, A. D’Agostino, C. Ottoni, M. Carra, S. Vukojičić, M. Constantinescu, D. Antonović, T. Douglas Price, D. Borić

**Affiliations:** DANTE - Diet and ANcient TEchnology Laboratory, Department of Oral and Maxillo-Facial Sciences, Sapienza University of Rome, Via Caserta 6, Rome 00161, Italy; BioArCh University of York, UK; Laboratory of General Botany, Department of Biology, University of Rome “Tor Vergata,” Via della Ricerca Scientifica 1, 00133 Rome, Italy; University of Belgrade, Faculty of Biology, Institute of Botany and Botanical Garden “Jevremovac”, Takovska 43, 11000 Belgrade, Serbia; Romanian Academy, Institute for Anthropological Research “Francisc I. Rainer”, Bucarest, Romania; Institute of Archaeology, Knez Mihailova 35/IV, 11000 Belgrade, Serbia; Department of Anthropology, University of Wisconsin, Madison, USA; The Italian Academy for Advanced Studies in America, Columbia University, 1161 Amsterdam Avenue, New York, NY 10024, USA; Department of Environmental Biology, Sapienza University of Rome, P.le A. Moro 5, Rome 00185, Italy

## Abstract

Forager focus on wild cereal plants has been documented in the core zone of domestication in southwestern Asia, while evidence for forager use of wild grass grains remains sporadic elsewhere. In this paper, we present starch grain and phytolith analyses of dental calculus from 61 Mesolithic and Early Neolithic individuals from five sites in the Danube Gorges of the central Balkans. This zone was inhabited by likely complex Holocene foragers for several millennia before the appearance of the first farmers ∼6200 cal BC. We also analyzed forager ground stone tools for evidence of plant processing. Our results based on the study of dental calculus show that certain species of Poaceae (species of the genus *Aegilops*) were used since the Early Mesolithic, while ground stone tools exhibit traces of a developed grass grain processing technology. The adoption of domesticated plants in this region after ∼6500 cal BC might have been eased by the existing familiarity with wild cereals.

## Introduction

Forager knowledge and consistent use of wild cereals are still debated and poorly documented outside of the assumed centers of domestication in southwestern Asia (Kotsakis, 2003). For some time, it has been claimed that in the Balkans some forms of intense gathering or incipient human management of local plants and animals might have occurred before the full-blown transition to the Neolithic (Clarke, 1978; Halstead, 1996; Kotsakis, 2003; Kotzamani and Livarda, 2014; Srejović, 1988; y’Edynak and Fleisch, 1983), partly due to the region’s geographical proximity to the Near East. However, the hypothesis of a systematic use of wild grasses of the Poaceae family (e.g., *Aegilops* spp.; *Hordeum* spp.) during the Mesolithic remains to be verified in this region.

In southeastern Europe, specifically in its Mediterranean zone, where one would expect a greater spectrum of small seeded grasses, fruits, and nuts, forager consumption of wild cereals is well documented only at Franchthi cave in Greece. Here, wild barley (*Hordeum* sp.) appears in the archaeobotanical record starting in the Late Upper Palaeolithic and throughout the Mesolithic, along with oat (*Avena* sp.), pulses (*Lens sp. Mill*.), bitter vetch (*Vicia ervilia* (L.) Willd.), almond (*Prunus amygdalus Batsch*), and terebinth (*Pistacia cf. lentiscus* L.)(Hansen, 1991; Van Andel et al., 1987). More recently, at Vlakno Cave in Croatia, starch granules of a wild species of barley (*Hordeum* spp.), along with those of oat (*Avena* spp.), were found in the dental calculus of a Mesolithic forager individual, dating to the late eight millennium cal BC burial (Cristiani et al., 2018).

Besides this type of evidence, data about the increase of cereal-type pollen in the Late Mesolithic come from palynological spectra from across Europe. Although the exclusive reliance on pollen evidence for inferring cultivation can be problematic, consistent evidence for interventions in the forest canopy, marked as disturbances in pollen spectra, might suggest anthropogenic activity. Due to low dispersal rates of cereal-type pollen grains as well as Cerealia-type pollens, their very presence in pollen spectra could be highly indicative of anthropic origin of disturbance phases (Edwards, 1989), and could be interpreted as forest clearances.

Recent methodological advances in our ability to analyse micro-residues in the form of micro-remains along with surface modifications and micro-residues on ground stone tools (henceforth GST) (Barton et al., 2018; Dubreuil and Nadel, 2015; Radini et al., 2017) have the potential to contribute to this old debate about Balkan and other prehistoric foragers’ familiarity with plant species. Moreover, far from seeing foragers as passive recipients of novelties arriving from Neolithic groups at the time of agricultural transitions, there is now growing evidence of the active role of hunter-gatherers in shaping their landscape ecologies, including plant management, and manipulation of ecosystems through niche constructing (Lombardo et al., 2020; Rowley-Conwy and Layton, 2011; Smith, 2011).

We examine these pertinent issues in hunter-gatherer research by studying dental remains and GSTs found at Mesolithic and Neolithic sites in the Danube Gorges area of the north-central Balkans between present-day Serbia and Romania (Figs. 1,2,3). This is one of the best researched areas of Europe regarding the Mesolithic-Neolithic transition period with more than 20 sites spanning the duration of the Epipalaeolithic through to the Mesolithic and Early Neolithic (∼13,000–5500 cal BC) (Bonsall, 2008; Borić, 2011; Radovanović, 1996; Srejović, 1972). Open-air sites began appearing in the archaeological record with the start of the Holocene warming on river terrace promontories in the vicinity of strong whirlpools, narrows, and rapids of the Danube, which facilitated intense fishing operations (Borić, 2011). The Early and Middle Mesolithic (∼9600–7300 cal BC) deposits at many sites are damaged by later Mesolithic and Neolithic intrusions, but a number of burials have directly been AMS-dated to these early phases. From the Early Mesolithic onwards, these sites became places for a continuous interment of the dead (Borić, 2016; Borić et al., 2014; Radovanović, 1996), thus creating a substantial mortuary record, which is in the excess of 500 individuals. Osteological collections allowed for a host of bioarchaeological analyzes to be applied on this material (Bonsall et al., 1997; Borić et al., 2004; Borić and Price, 2013; Mathieson et al., 2018). Fishing seems to have remained one of the main subsistence foci throughout the Holocene, with a possible intensification during the Late Mesolithic (∼7300–6200 cal BC), during which period there is an intense inhabitation of the area, with recognizable features in the archaeological record, such as stone-lined rectangular hearths and abundant primary burials placed as extended inhumations parallel to the Danube River. Between ∼6200 and 5900 cal BC, there are clearest indications based on both material culture associations and isotope and genomic data (Borić and Price, 2013; Mathieson et al., 2018) that the local Mesolithic foragers came into contact with the first Neolithic groups appearing in this region, and who likely founded several new sites in this area, especially in the downstream part of the region. These documented encounters of two different cultural groups are most clearly observed at the site of Lepenski Vir (Boric, 2016; Borić et al., 2018). After ∼5900 cal BC, it seems that the forager cultural specificity was lost and that various sites remained to be used as typical Early Neolithic Starčevo culture villages up until ∼5500 cal BC, when most of the previously used locales were abandoned.

**Fig. 1.**
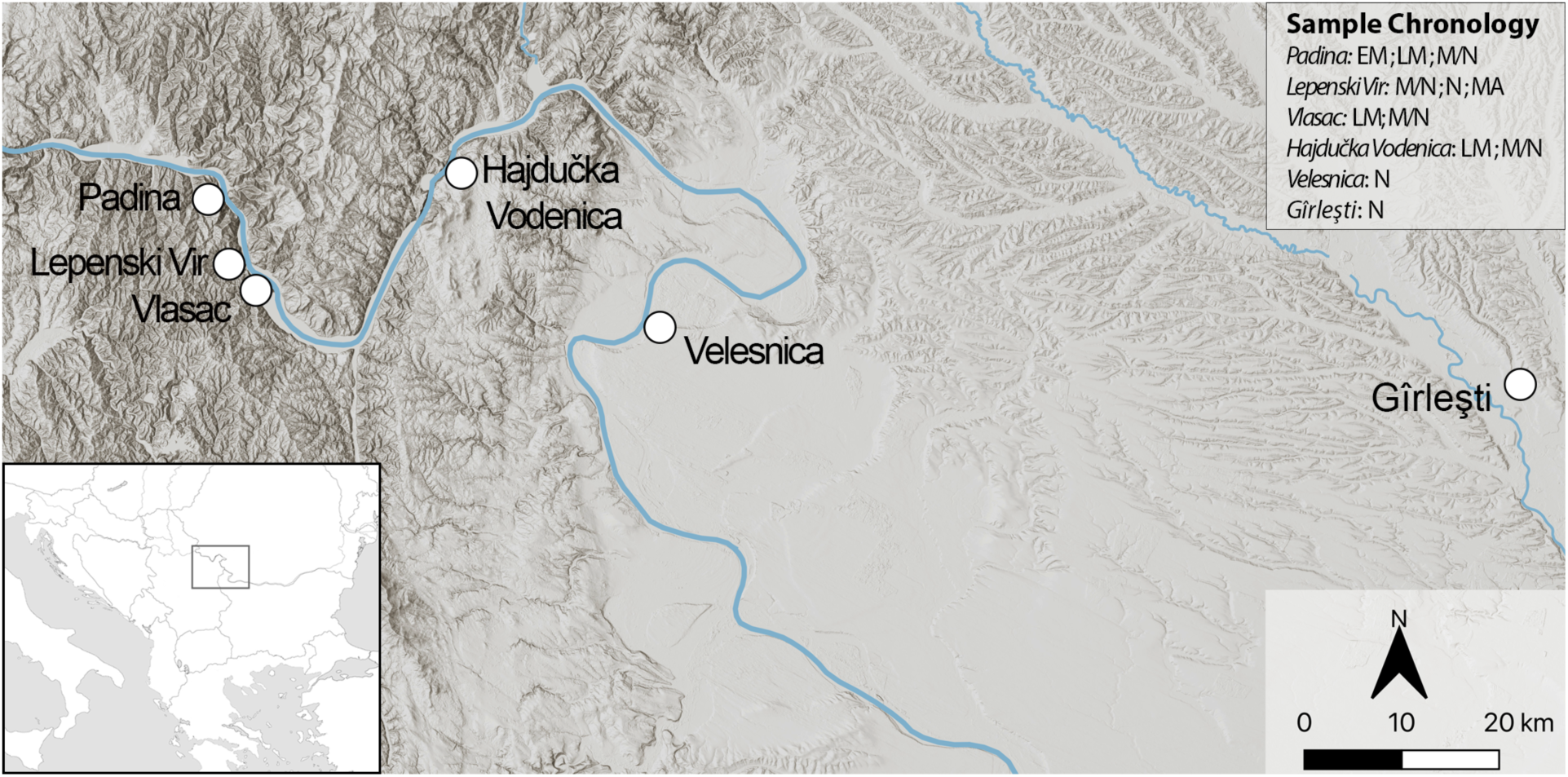
Sites in the central Balkans investigated in the article, which provided dental calculus and ground stone tools. EM = Early Mesolithic; LM = Late Mesolithic; M/N= Mesolithic/Neolithic; N = Neolithic.

**Fig. 2.**
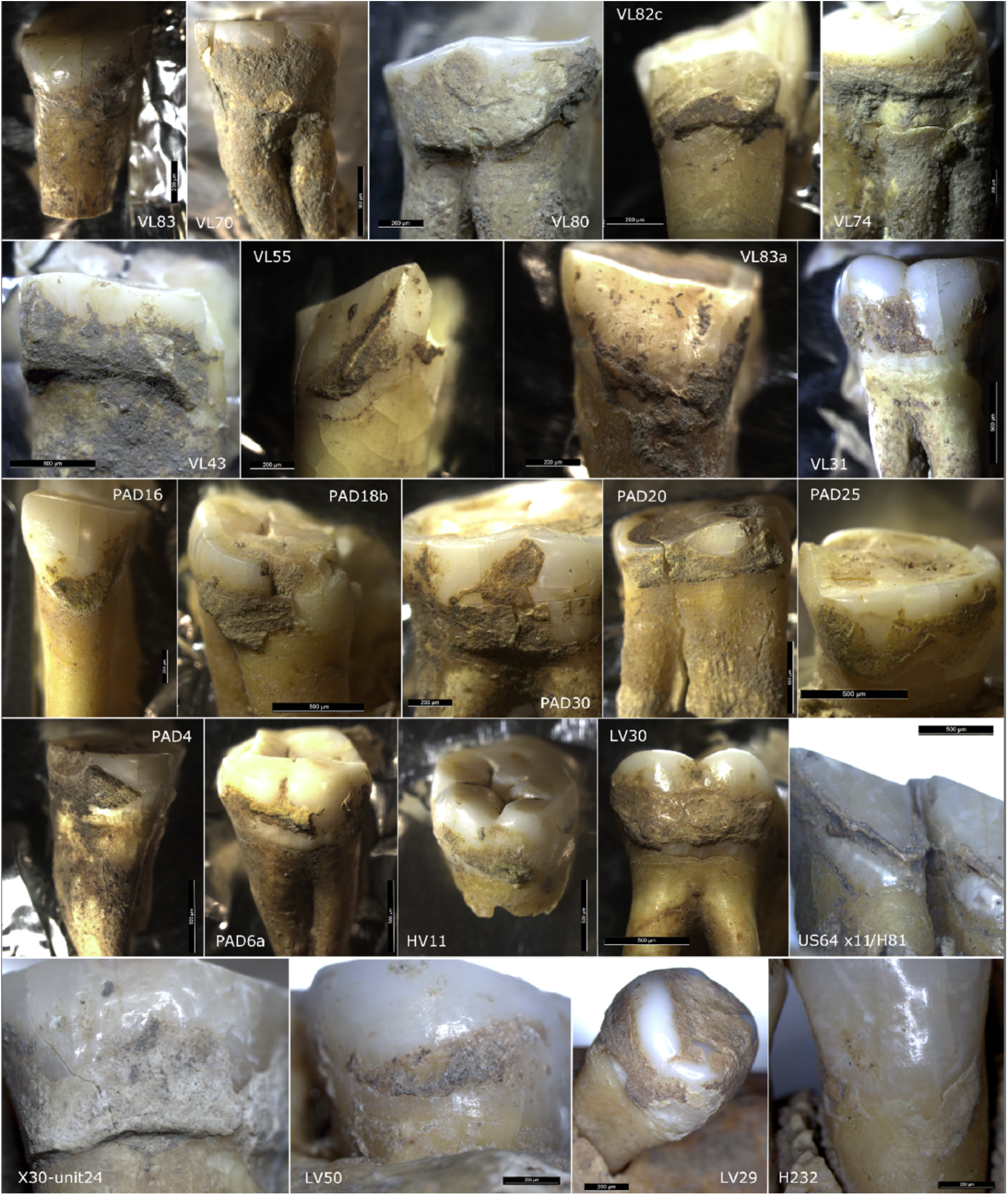
Studied teeth photographed under the microscope before dental calculus sampling.

**Fig. 3.**
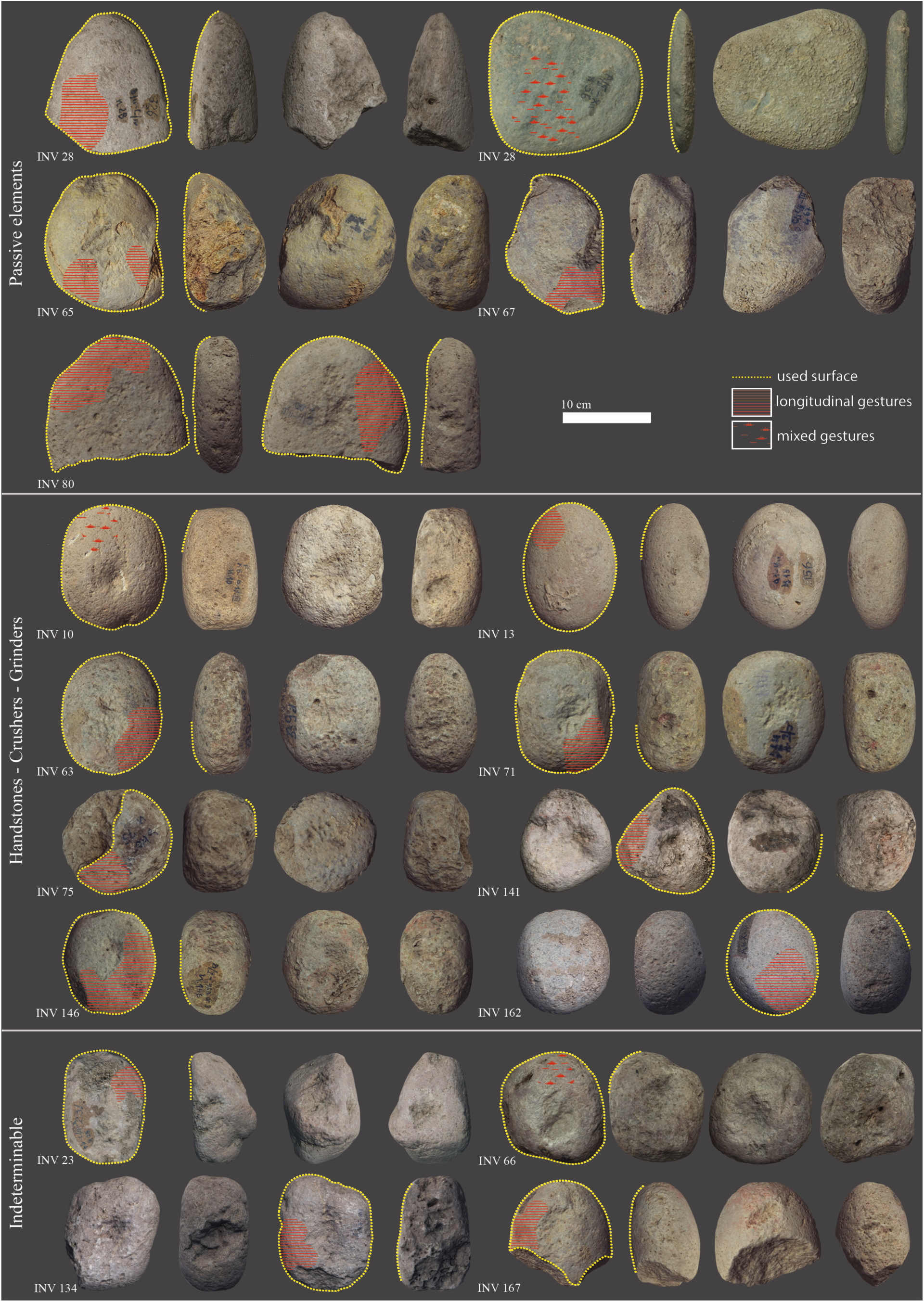
Late Mesolithic GSTs from the site of Vlasac featuring use-wear traces and residues related to plant food processing.

While sources of animal protein in the diet of Mesolithic-Neolithic inhabitants of the area are well understood by now, the significance of plant foods in this region has remained less well known. Non-flaked tools such as pestles, grinders, crushers, and anvils have recently been associated with fruit, seed, and nut processing in early prehistoric and ethnographic contexts (de Beaune, 2004; Dubreuil and Nadel, 2015; Hamon et al., 2021; Pardoe et al., 2019; Wright, 1994). However, this category of artefacts is primarily documented from the Late Mesolithic onwards and only sporadically associated with earlier periods in the region of the Danube Gorges (Antonović et al., 2006; Borić et al., 2014; Srejović and Letica, 1978).

Such a lack of evidence about the role of plant foods in Mesolithic stems from very limited attempts to recover macro-botanical remains through intense sediment flotation, which has only been applied at two more recently excavated sites in this region—Schela Cladovei (Mason et al., 1996) and Vlasac (Borić et al., 2014; Mason et al., 1996). Despite a generally poor preservation of plant remains due to taphonomic issues, recent carpological analyses indicated a relatively wide spectrum of wild resources available to the local Mesolithic foragers. These included drupes, fruits, and berries (Marinova et al., 2013), among which cornelian cherry (*Cornus mas*), hazelnut (*Corylus avellana*), and elderberry (*Sambucus nigra*) were the most frequent taxa (Filipović et al., 2010; Marinova et al., 2013). Molecular record of *C. avellana* and *S. nigra* was also found preserved in the dental calculus of two Late Mesolithic individuals from Vlasac, which underwent metagenomic analysis (Ottoni et al., 2021). Moreover, in the study region of the Danube Gorges, at the site of Vlasac, presumed human palaeofeces contained pollen of Amaranthaceae and Cerealia (Cârciumaru, 1978). Evidence from the Mesolithic levels of the site of Icoana, located in the same region, has suggested local cereal cultivation (Cârciumaru, 1973). Pollen provides only indirect evidence for consumption and cultivation and, unfortunately, the 1960–70s excavations of the sites in the Danube Gorges did not involve any flotation of contextual units associated with burning in domestic contexts. While the extensive program of flotation at the site of Vlasac in the course of more recent work (2006–2009) has not led to the discovery of macro-botanical remains of wild or domesticated cereal grains, it should also be emphasized that the new excavations at this site have taken place in a marginal, upslope part of the site with little or no evidence of domestic features associated with burning that might have preserved macro-botanical remains (Borić et al., 2014). More recently, starch granules identified in dental calculus within a sample of 12 individuals provided evidence for the consumption of domesticated cereals at the site of Vlasac during the Late Mesolithic (Cristiani et al., 2016).

Hence, plant debris recovered in human dental calculus constitute the most reliable line of evidence to unveil the role of plants in the local forager diets. Our previous pilot study has provided the first evidence of domesticated cereal grains and plant food consumption in the Late Mesolithic from the analyzes of dental calculus (Cristiani et al., 2016). Based on a more robust sample of human dental calculus, which now also involves numerous Early Mesolithic individuals not included in our previous study, and complementary functional evidence from the most conspicuous assemblage of Mesolithic GSTs from the Danube Gorges, the present study details forager use of certain species of the Triticeae tribe and other plant foods in the region already since ∼9500 cal BC.

## Results

### Dental calculus

Starch granules were almost ubiquitous in the analyzed individuals and many of them were found still in part associated with dental calculus remains. Six morphotypes have been retrieved in this study (Tables 1,2). We have not attempted the identification of starch granules less than 5 μm to avoid misinterpretation of transitory and small storage starch granules (Haslam, 2004).

**Table 1.** Details of dental calculus sampled for the study (n=61).

| Site | Lab label | Cultural attribution | AMS dates |  |  |  | Calculus location |  |  |
| --- | --- | --- | --- | --- | --- | --- | --- | --- | --- |
|  |  |  | Lab Code | 14C age (BP) | Reservoir effect corrected age (BP) | 95.4% confidence, cal BC | Tooth | Surface | Weight (mg) |
| Padina | PAD20 | Early Meso |  |  |  |  | 17 | buccal | 9.6 |
| Padina | PAD25 | Early Meso |  |  |  |  | 38 | buccal | 9.59 |
| Padina | PAD15 | Early Meso | OxA-17145 | 9310±44 | 8870±63 | 8237–7761 (Boric, 2011) | 38 | lingual | 9.58 |
| Padina | PAD16 a | Early Meso | PSU-2407 | 9340±35 | 8907±66 | 8265–7820 (Mathieson et al., 2018) | 34 | buccal | 9.62 |
| Padina | PAD18 b | Early Meso | PSU-2376 | 9715±40 | 9424±55 | 9115–8555 (Mathieson et al., 2018) | 48 | lingual | 9.67 |
| Padina | PAD9 | Early Meso | AA-57771 | 9920±100 | 9480±110 | 9221–8548 (Boric, 2011) | 42, 46 | lingual | 9.59 |
| Padina | PAD11 | Early Meso | OxA-16938 | 9665±54 | 9225±70 | 8616–8296 (Boric, 2011) | 27 | lingual | 9.57 |
| Padina | PAD12 | Early Meso | BM-1146 | 9331±58 | – | 8753–8351 (Boric, 2011) | 27 | lingual | 9.76 |
| Padina | PAD17 | Early Meso | PSU-2375 | 9505±35 | 9105±62 | 8535–8230 (Mathieson et al., 2018) | 25 | buccal | 9.64 |
| Lepenski Vir | LV50 | Early Meso | BA-10651 | 9455±38 | 9082±62 | 8532–8208 (Borić, 2019) | 35 | buccal | 9.99 |
| Lepenski Vir | LV20 | Early Meso | OxA-39629 | 10,268±38 | 9928±58 | 9665–9275 (this paper) | 48 | lingual | 9.73 |
| Padina | PAD26 | Early Meso |  |  |  |  | 14 | buccal | 9.58 |
| Padina | PAD6 | Early Meso |  |  |  |  | 47 | lingual | 9.65 |
| Padina | PAD2 | Late Meso | BM-1143 | 7738±51 | – | 6648–6470 (Boric, 2011) | 36 | lingual | 9.68 |
| Hajdučka Vodenica | HV25/2 6 | Late Meso |  |  |  |  | 44 | buccal | 9.60 |
| Hajdučka Vodenica | HV29 | Late Meso | AA-57774 | 8151±60 | 7711±75 | 6680–6434 (Boric, 2011) | 48 | lingual | 10.72 |
| Hajdučka Vodenica | HV8 | Late Meso | OxA-13613 | 8456±37 | 8016±58 | 7076–6699 (Boric, 2011) | 48 | buccal | 9.61 |
| Hajdučka Vodenica | HV11 | Late Meso |  |  |  |  | 48 | buccal | 9.71 |
| Hajdučka Vodenica | HV profil A | Late Meso |  |  |  |  | 27 | buccal | 9.70 |
| Hajdučka Vodenica | HV30 | Late Meso |  |  |  |  | 27 | buccal | 9.56 |
| Vlasac | VL82c | Late Meso |  |  |  |  | 42 | buccal | 9.68 |
| Vlasac | VL2 | Late Meso |  |  |  |  | 14 | buccal | 9.54 |
| Vlasac | VL80a | Late Meso |  |  |  |  | 26 | lingual | 9.84 |
| Vlasac | VL55 | Late Meso |  |  |  |  | 33 | lingual | 9.64 |
| Vlasac | VL74 | Late Meso |  |  |  |  | 28 | lingual | 9.70 |
| Vlasac | VL83 | Late Meso | OxA-5826 | 8200±90 | 7760±100 | 7024–6430 (Boric, 2011) | 24 | lingual | 9.62 |
| Vlasac | VL43 | Late Meso |  |  |  |  | 27 | lingual |  |
| Vlasac | VL31 | Late Meso | AA-57777 | 8196±69 | 7756±82 | 6823–6436 (Boric, 2011) | 26 | buccal | 9.58 |
| Vlasac | VL45 | Late Meso | AA-57778 | 8117±62 | 7677±77 | 6654–6411 (Boric et al., 2008) | 38 | buccal | 9.50 |
| Vlasac | VL70 | Late Meso |  |  |  |  | 17 | buccal | 10.42 |
| Vlasac | VL79 | Late Meso |  |  |  |  | 16 | buccal | 9.60 |
| Vlasac | U44 | Late Meso |  |  |  |  | 27 | buccal | 9.96 |
| Vlasac | H232 | Late Meso | OxA-20702 | 7725±40 |  | 6620–6480 (Boric, 2011) | 28 | lingual | 9.92 |
| Vlasac | H317 | Late Meso | PSU-2381 | 8110±35 | 7625±71 | 6635–6375 (Mathieson et al., 2018) | 26, 36 | lingual | 9.73 |
| Vlasac | U115 | Late Meso |  |  |  |  | 28 | buccal | 9.95 |
| Vlasac | U327 | Late Meso | PSU-2382 | 8045±30 | 7728±51 | 6645–6465 (Mathieson et al., 2018) | 17 | buccal | 9.94 |
| Vlasac | U64 x.11/H8 1 | Late Meso | OxA-20762 | 8125±45 | 7685 ± 64 | 6590–6470 (Boric, 2011) | 20, 26, 27, 29, 30, 31 | lingual | 9.92 |
| Vlasac | H341 | Late Meso |  |  |  |  | 1 | buccal | 10.12 |
| Vlasac | VL48 | Late Meso |  |  |  |  | 34 | lingual | 10.06 |
| Vlasac | U326 | Late Meso |  |  |  |  | 1, 2 | buccal | 9.1 |
| Vlasac | H341 | Late Meso |  |  |  |  |  |  |  |
| Vlasac | U222 x.18 | Late Meso |  |  |  |  | 2 | buccal | 9.54 |
| Lepenski Vir | LV28 | Meso-Neo |  |  |  | – | 43 | buccal | 9.58 |
| Lepenski Vir | LV79a | Meso-Neo | OxA-25091 | 7605±38 | 7119±74 | 6206–5840 (Bonsall et al., 2015) | 33 | buccal | 9.69 |
| Hajdučka Vodenica | HV16 | Meso-Neo |  |  |  |  | 36 | lingual | 9.54 |
| Hajdučka Vodenica | HV19 | Meso-Neo |  |  |  |  | 37 | buccal | 9.58 |
| Hajdučka Vodenica | HV13 | Meso-Neo | AA-57773 | 7435±70 | 6995±83 | 6016–5726 (Boric, 2011) | 17 | lingual | 9.56 |
| Padina | PAD4 | Meso-Neo | AA-57769 | 7518±72 | 7078±85 | 6061–5841 (Ottoni et al., 2021) | 48 | buccal | 9.73 |
| Padina | PAD5 | Meso-Neo | AA-57770 | 7598±72 | 7158±85 | 6224–5878 (Boric, 2011) | 15 | buccal | 8.10 |
| Vlasac | U24 x.30 | Meso-Neo |  |  |  |  | 32 | lingual | 9.97 |
| Vlasac | H53 | Meso-Neo | OxA-16544 | 7035±40 | – | 6006–5838 | 3, 28, 29 | lingual | 10.04 |
| Lepenski Vir | LV4 | E. Neolithic |  |  |  |  | 33 | buccal | 9.64 |
| Lepenski Vir | LV73 | E. Neolithic | BA-10652 | 7265±30 | 6973±48 | 5981–5742 (Borić, 2019) | 34 | buccal | 9.69 |
| Lepenski Vir | LV8 | E. Neolithic | AA-58319<br>OxA-25207 | 6825±51<br>7097±36 | 6690±54<br>6984±39 | 5715–5521<br>5981–5751 (Borić, 2019) | 44 | lingual | 9.69 |
| Lepenski Vir | LV32A | E. Neolithic | OxA-5828 | 7270±90 | 7032±95 | 6066–5727 (Bonsall et al., 1997) | 42, 43, 36 | buccal | 9.77 |
| Lepenski Vir | LV17 | E. Neolithic | AA-58320 | 7007±48 | 6787±53 | 5776–5575 (Bonsall et al., 2015) | 15 | lingual | 9.10 |
| Padina | PAD30 | Bronze Age | PSU-2379 |  |  |  | 47 | buccal | 9.69 |
| Velesnica | 2A | Neo | OxA-19191 | 7409±38 | 7196±47 | 6210–5990 (Bonsall et al., 2015) | 8 | lingual | 9.74 |
| Velesnica | 2D | E. Neolithic | OxA-19210 | 7327±38 | 7183±42 | 6205–5985 (Bonsall et al., 2015) | 9 | lingual | 7.2 |
| Girlestj |  | E. Neolithic |  |  |  |  | 2 | lingual | 8.22 |
| Lepenski Vir | LV30 | Medieval | OxA-25218 | 427±23 | 281±30 | AD1492–1648 (not corrected) (Bonsall et al., 2015) | 16 | lingual | 9.67 |

**Table 2.**
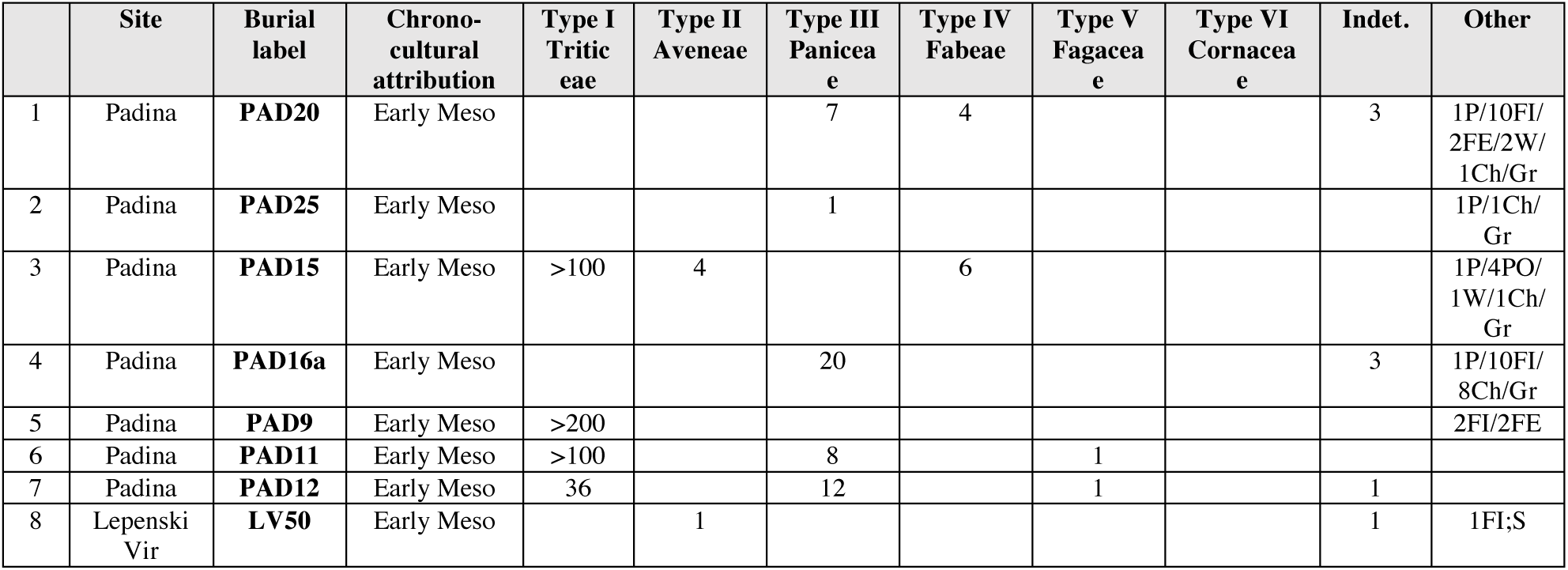
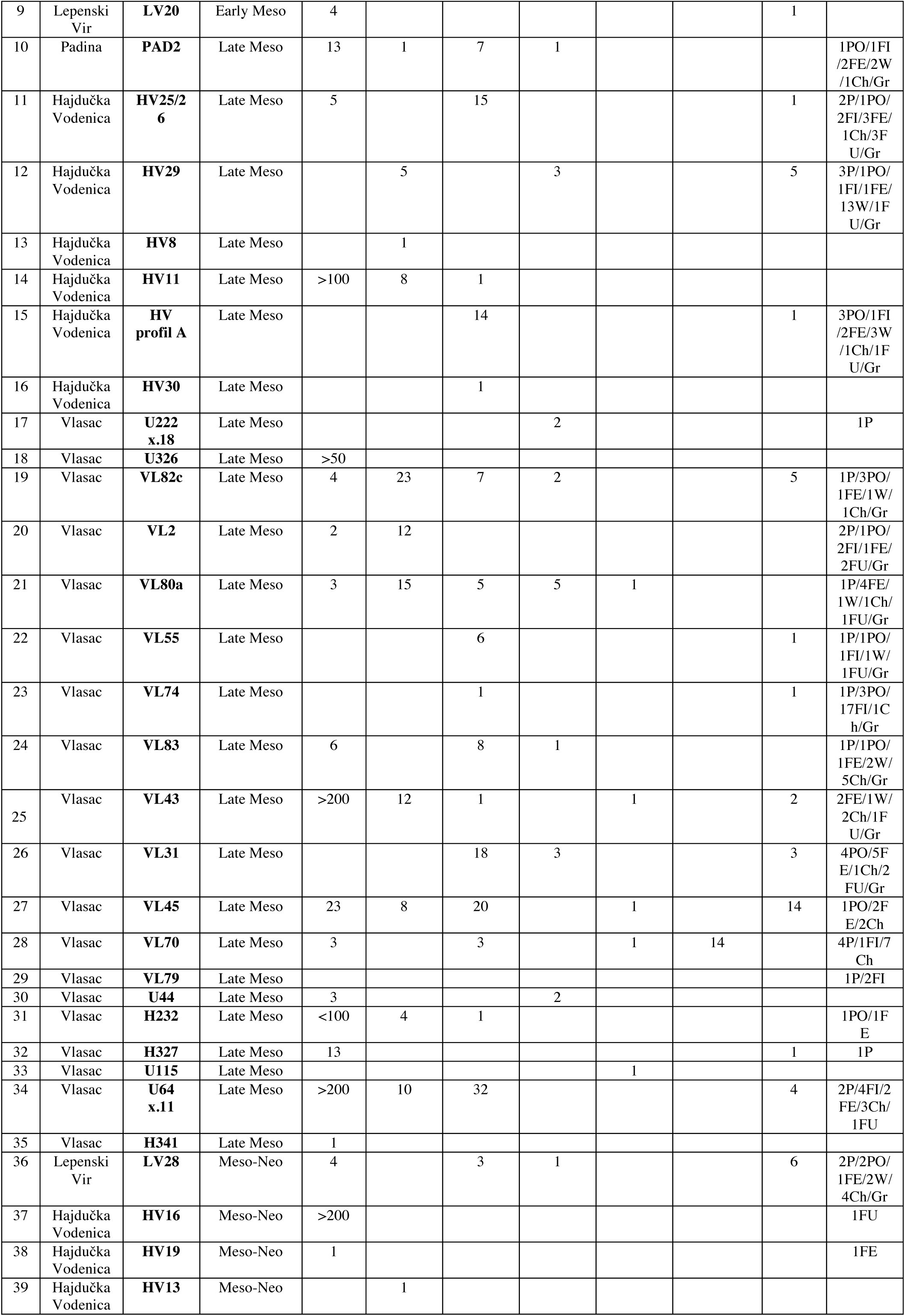
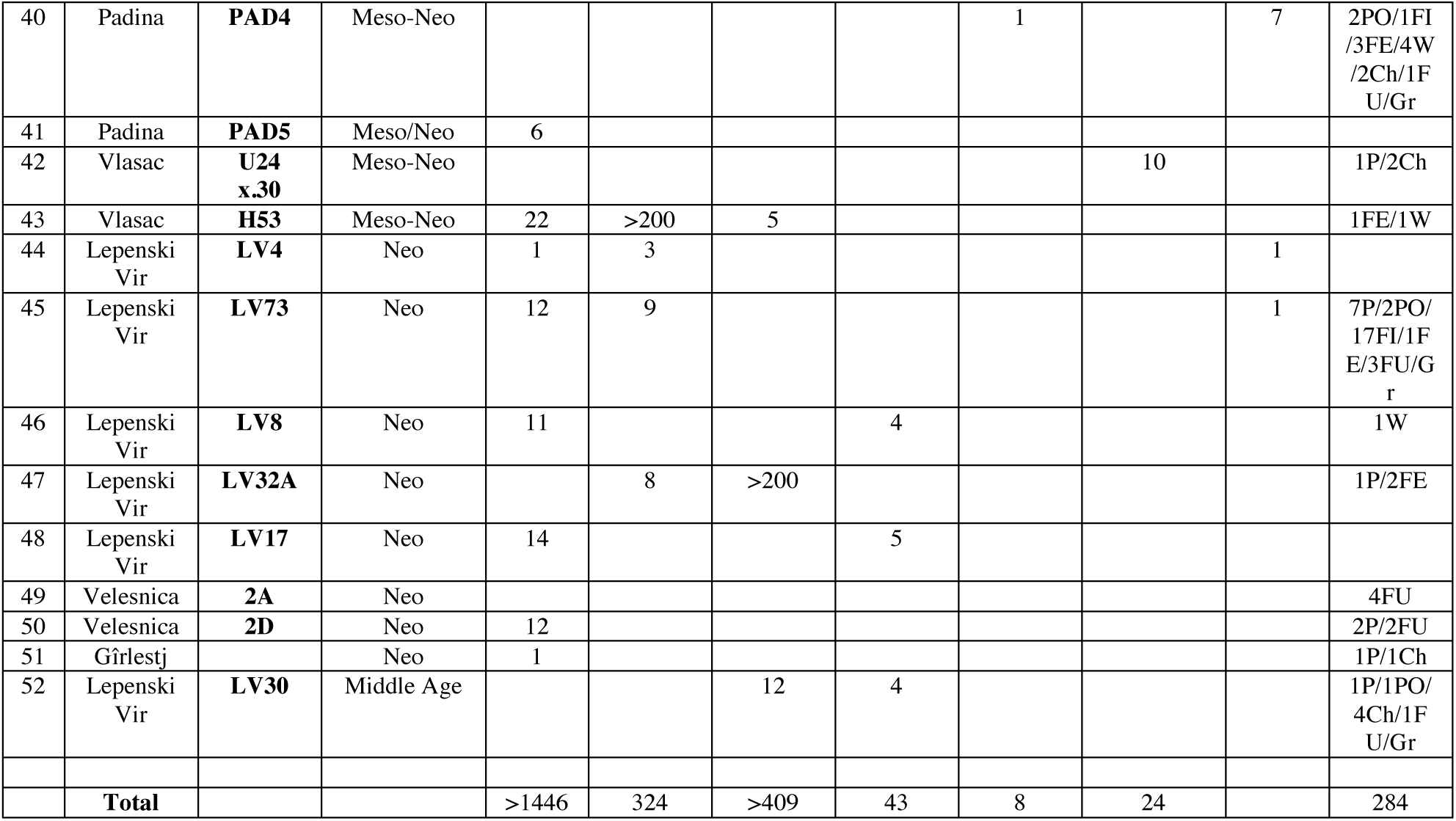
Details of the micro-debris (starch granules and other micro-remains) found in the archaeological dental calculus samples (PO=Pollen; W=Wood; Ch=Charcoal; Gr=Grit; P=Phytoliths; FE=Feathers; FI=Fibers; FU=Fungi; S=Smoke) (n=52).

|  | Site | Burial label | Chrono-cultural attribution | Type I Tritic eae | Type II Aveneae | Type III Panicea e | Type IV Fabeae | Type V Fagacea e | Type VI Cornacea e | Indet. | Other |
| --- | --- | --- | --- | --- | --- | --- | --- | --- | --- | --- | --- |
| 1 | Padina | PAD20 | Early Meso |  |  | 7 | 4 |  |  | 3 | 1P/10FI/<br>2FE/2W/<br>1Ch/Gr |
| 2 | Padina | PAD25 | Early Meso |  |  | 1 |  |  |  |  | 1P/1Ch/<br>Gr |
| 3 | Padina | PAD15 | Early Meso | >100 | 4 |  | 6 |  |  |  | 1P/4PO/<br>1W/1Ch/<br>Gr |
| 4 | Padina | PAD16a | Early Meso |  |  | 20 |  |  |  | 3 | 1P/10FI/<br>8Ch/Gr |
| 5 | Padina | PAD9 | Early Meso | >200 |  |  |  |  |  |  | 2FI/2FE |
| 6 | Padina | PAD11 | Early Meso | >100 |  | 8 |  | 1 |  |  |  |
| 7 | Padina | PAD12 | Early Meso | 36 |  | 12 |  | 1 |  | 1 |  |
| 8 | Lepenski Vir | LV50 | Early Meso |  | 1 |  |  |  |  | 1 | 1FI;S |
| 9 | Lepenski Vir | **LV20** | Early Meso | 4 |  |  |  |  |  | 1 |  |
| 10 | Padina | **PAD2** | Late Meso | 13 | 1 | 7 | 1 |  |  |  | 1PO/1FI/2FE/2W/1Ch/Gr |
| 11 | Hajdučka Vodenica | **HV25/26** | Late Meso | 5 |  | 15 |  |  |  | 1 | 2P/1PO/2FI/3FE/1Ch/3FU/Gr |
| 12 | Hajdučka Vodenica | **HV29** | Late Meso |  | 5 |  | 3 |  |  | 5 | 3P/1PO/1FI/1FE/13W/1FU/Gr |
| 13 | Hajdučka Vodenica | **HV8** | Late Meso |  | 1 |  |  |  |  |  |  |
| 14 | Hajdučka Vodenica | **HV11** | Late Meso | >100 | 8 | 1 |  |  |  |  |  |
| 15 | Hajdučka Vodenica | **HV profil A** | Late Meso |  |  | 14 |  |  |  | 1 | 3PO/1FI/2FE/3W/1Ch/1FU/Gr |
| 16 | Hajdučka Vodenica | **HV30** | Late Meso |  |  | 1 |  |  |  |  |  |
| 17 | Vlasac | **U222 x.18** | Late Meso |  |  |  | 2 |  |  |  | 1P |
| 18 | Vlasac | **U326** | Late Meso | >50 |  |  |  |  |  |  |  |
| 19 | Vlasac | **VL82c** | Late Meso | 4 | 23 | 7 | 2 |  |  | 5 | 1P/3PO/1FE/1W/1Ch/Gr |
| 20 | Vlasac | **VL2** | Late Meso | 2 | 12 |  |  |  |  |  | 2P/1PO/2FI/1FE/2FU/Gr |
| 21 | Vlasac | **VL80a** | Late Meso | 3 | 15 | 5 | 5 | 1 |  |  | 1P/4FE/1W/1Ch/1FU/Gr |
| 22 | Vlasac | **VL55** | Late Meso |  |  | 6 |  |  |  | 1 | 1P/1PO/1FI/1W/1FU/Gr |
| 23 | Vlasac | **VL74** | Late Meso |  |  | 1 |  |  |  | 1 | 1P/3PO/17FI/1Ch/Gr |
| 24 | Vlasac | **VL83** | Late Meso | 6 |  | 8 | 1 |  |  |  | 1P/1PO/1FE/2W/5Ch/Gr |
| 25 | Vlasac | **VL43** | Late Meso | >200 | 12 | 1 |  | 1 |  | 2 | 2FE/1W/2Ch/1FU/Gr |
| 26 | Vlasac | **VL31** | Late Meso |  |  | 18 | 3 |  |  | 3 | 4PO/5FE/1Ch/2FU/Gr |
| 27 | Vlasac | **VL45** | Late Meso | 23 | 8 | 20 |  | 1 |  | 14 | 1PO/2FE/2Ch |
| 28 | Vlasac | **VL70** | Late Meso | 3 |  | 3 |  | 1 | 14 |  | 4P/1FI/7Ch |
| 29 | Vlasac | **VL79** | Late Meso |  |  |  |  |  |  |  | 1P/2FI |
| 30 | Vlasac | **U44** | Late Meso | 3 |  |  | 2 |  |  |  |  |
| 31 | Vlasac | **H232** | Late Meso | <100 | 4 | 1 |  |  |  |  | 1PO/1FE |
| 32 | Vlasac | **H327** | Late Meso | 13 |  |  |  |  |  | 1 | 1P |
| 33 | Vlasac | **U115** | Late Meso |  |  |  |  | 1 |  |  |  |
| 34 | Vlasac | **U64 x.11** | Late Meso | >200 | 10 | 32 |  |  |  | 4 | 2P/4FI/2FE/3Ch/1FU |
| 35 | Vlasac | **H341** | Late Meso | 1 |  |  |  |  |  |  |  |
| 36 | Lepenski Vir | **LV28** | Meso-Neo | 4 |  | 3 | 1 |  |  | 6 | 2P/2PO/1FE/2W/4Ch/Gr |
| 37 | Hajdučka Vodenica | **HV16** | Meso-Neo | >200 |  |  |  |  |  |  | 1FU |
| 38 | Hajdučka Vodenica | **HV19** | Meso-Neo | 1 |  |  |  |  |  |  | 1FE |
| 39 | Hajdučka Vodenica | **HV13** | Meso-Neo |  | 1 |  |  |  |  |  |  |
| 40 | Padina | **PAD4** | Meso-Neo |  |  |  |  | 1 |  | 7 | 2PO/1FI<br>/3FE/4W<br>/2Ch/1F<br>U/Gr |
| 41 | Padina | **PAD5** | Meso/Neo | 6 |  |  |  |  |  |  |  |
| 42 | Vlasac | **U24**<br>**x.30** | Meso-Neo |  |  |  |  |  | 10 |  | 1P/2Ch |
| 43 | Vlasac | **H53** | Meso-Neo | 22 | >200 | 5 |  |  |  |  | 1FE/1W |
| 44 | Lepenski<br>Vir | **LV4** | Neo | 1 | 3 |  |  |  |  | 1 |  |
| 45 | Lepenski<br>Vir | **LV73** | Neo | 12 | 9 |  |  |  |  | 1 | 7P/2PO/<br>17FI/1F<br>E/3FU/G<br>r |
| 46 | Lepenski<br>Vir | **LV8** | Neo | 11 |  |  | 4 |  |  |  | 1W |
| 47 | Lepenski<br>Vir | **LV32A** | Neo |  | 8 | >200 |  |  |  |  | 1P/2FE |
| 48 | Lepenski<br>Vir | **LV17** | Neo | 14 |  |  | 5 |  |  |  |  |
| 49 | Velesnica | **2A** | Neo |  |  |  |  |  |  |  | 4FU |
| 50 | Velesnica | **2D** | Neo | 12 |  |  |  |  |  |  | 2P/2FU |
| 51 | Gîrlestj |  | Neo | 1 |  |  |  |  |  |  | 1P/1Ch |
| 52 | Lepenski<br>Vir | **LV30** | Middle Age |  |  | 12 | 4 |  |  |  | 1P/1PO/<br>4Ch/1F<br>U/Gr |
|  | **Total** |  |  | >1446 | 324 | >409 | 43 | 8 | 24 |  | 284 |

**Table 3.**
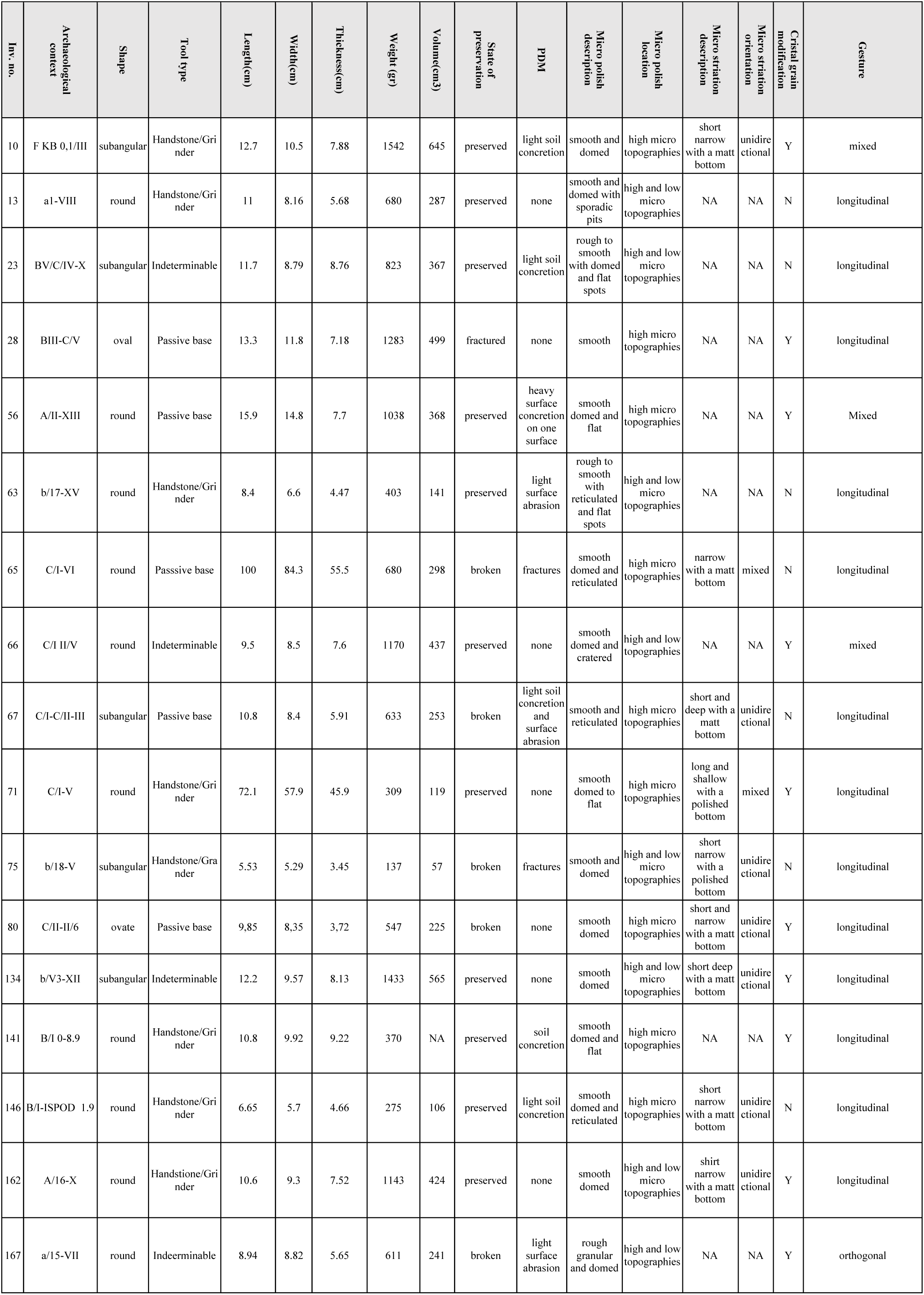
Late Mesolithic GSTs from the site of Vlasac.

#### Type I

Size, shape, morphology and bimodal distribution that characterize granules of this type are encountered in Europe only in the members of the plant tribe Triticeae (Poaceae family) and considered diagnostic features for taxonomic identification (Henry and Piperno, 2008; Stoddard, 1999; Yang and Perry, 2013). Such distribution involves the presence of large granules (A-Type), mostly with a clear, round to sub-oval in 2D shape, ranging between 21.1 and 62.7 μm in maximum dimensions (mean size of 41.9 μm), lenticular 3D shape with equatorial groove always visible, a central hilum and high density of deep lamellae concentrated in the mesial part; and small granules (B-Type) with round/sub-oval shapes, a central hilum, generally smaller than 10 μm (Geera et al., 2006; Stoddard, 1999; Yang and Perry, 2013). A-Type granules possess diagnostic features while smaller B-Type granules are rarely diagnostic to taxa (Yang and Perry, 2013). However, in our archaeological population, variability in the proportion and dimension of small B-type granules has been noticed, resulting in a unimodal granule size distribution without a clear distinction between A and B granules in some cases. Several studies (Howard et al., 2011; Stoddard and Sarker, 2000) suggested that this characteristic is common in the species of the genus *Aegilops* of the Triticeae tribe and can be attributed to both environment (Blumenthal et al., 1995, 1994) and genetics (Stoddard and Sarker, 2000). A unimodal starch granule size distribution characterized by normal A-type granules and a lack/reduced quantity of B-type granules was also evident in our modern reference collection of local *Aegilops* species (Fig.6). Based on these observations, Type I category was further divided into two subtypes (Ia and Ib). In subtype Ia, B-Type granules are small, dimensionally uniform (up to 12 μm) and round in shape (fig.6). Conversely, subtype Ib is characterized by a high variation in starch granule size not allowing for a distinction between A and B-type granule, resulting in a unimodal distribution (fig.9).

Type I (Ia and Ib) is very common in the analyzed samples (Table 2), as already emphasized in our earlier study albeit in different quantities (*24*). These starch granules were documented, often lodged in the amyloplast, in most of the analyzed Mesolithic individuals (5 for EM, 16 for LM, 5 for M/N), and in five EN individuals of our population (Table 2). A-Type granules recovered in EM and most of the LM individuals were very large (sometimes up to 35 μm), mostly with a clear, round shape, central hilum, and high density of deep lamellae mainly concentrated in the mesial part of the granules. Based on literature (Henry et al., 2011; Yang and Perry, 2013) and our extensive experimental and statistical results on modern botanical collection (Tables 4,5; Figs. 6, 7, 9), we confirm that these characteristics are consistent with A-Type granules of most species of the Triticeae tribe.

**Table 4.** Summary statistics of the length of wild grass grains and domestic cereal starch granules. **Table 4-source data 1**. Summary statistics of the length of wild grass grains and domestic cereal starch granules.

| Species | Min. | Max. | Mean | Median | StDev. | Range | IQR |
| --- | --- | --- | --- | --- | --- | --- | --- |
| *A. caudata* | 5.29 | 59.3 | 21.6 | 16.7 | 15.17 | 5.29 – 59.33 | 26.55 |
| *A. comosa* | 7.95 | 34.5 | 21.5 | 21.7 | 9.78 | 7.95 – 34.54 | 20.09 |
| *A. crassa* | 13.38 | 53.7 | 35.3 | 33.7 | 11.09 | 13.38 – 53.69 | 19.08 |
| *A. cylindrica* | 8.52 | 54.0 | 24.2 | 23.7 | 13.07 | 8.52 – 54.05 | 21.6 |
| *A. geniculata* | 11.61 | 47.0 | 26.3 | 26.0 | 8.39 | 11.61 – 47.03 | 12.87 |
| *A. neglecta recta* | 10.54 | 62.7 | 35.0 | 36.2 | 14.46 | 10.54 – 62.71 | 26.5 |
| *A. peregrina* | 9.84 | 53.6 | 27.8 | 25.9 | 9.89 | 9.84 – 53.62 | 11.34 |
| *A. speltoides tauschii* | 13.25 | 40.0 | 23.5 | 22.2 | 5.93 | 13.25 – 39.97 | 8.39 |
| *A. triuncialis* | 5.60 | 50.1 | 28.2 | 28.2 | 11.24 | 5.60 – 50.06 | 15.18 |
| *A. uniaristata* | 14.35 | 62.4 | 38.2 | 39.3 | 12.87 | 14.35 – 62.38 | 22.83 |
| *A. ventricosa* | 14.10 | 40.0 | 26.3 | 25.7 | 7.44 | 14.10 – 40.04 | 12.77 |
| *H. vulgare distichon* | 5.19 | 29.6 | 19.7 | 22.2 | 8.12 | 5.19 – 29.59 | 8.32 |
| *T. dicocum* | 6.17 | 41.5 | 16.5 | 12.8 | 8.66 | 6.17 – 41.55 | 14.07 |
| *T. monococum* | 6.68 | 36.6 | 20.1 | 19.1 | 7.11 | 6.68 – 36.61 | 10.44 |

**Table 5.** Pairwise Wilcoxon test performed on the length distribution of modern starches from *Aegilops, Hordeum*, and *Triticum* species (*p*-value: not significant/n.s.>0.05; ^*^<0.05; ^**^<0.01; ^***^<0.001). **Table 5-source data 2**. Length of modern starch granules of *Aegilops, Hordeum*, and *Triticum species*

|  | *A. caudat*<br>*a* | *A. comos*<br>*a* | *A. crass*<br>*a* | *A. cylindri*<br>*ca* | *A. genicul*<br>*ata* | *A. neglecta*<br>*recta* | *A. peregr*<br>*i-* | *A. speltoides*<br>*tauschii* | *A. triuncia*<br>*lis* | *A. uniarist*<br>*ata* | *A. ventrico*<br>*sa* | *H. vulgare*<br>*distichon* | *T. dicocc*<br>*um* |
| --- | --- | --- | --- | --- | --- | --- | --- | --- | --- | --- | --- | --- | --- |
| *A. comosa* | ns | - | - | - | - | - | - | - | - | - | - | - | - |
| *A. crassa* | *** | *** | - | - | - | - | - | - | - | - | - | - | - |
| *A. cylindrica* | ns | ns | *** | - | - | - | - | - | - | - | - | - | - |
| *A. geniculata* | ns | * | *** | ns | - | - | - | - | - | - | - | - | - |
| *A. neglecta recta* | *** | *** | ns | *** | *** | - | - | - | - | - | - | - | - |
| *A. peregrina* | * | ** | ** | ns | ns | ** | - | - | - | - | - | - | - |
| *A. speltoides tauschii* | ns | ns | *** | ns | ns | *** | * | - | - | - | - | - | - |
| *A. triuncialis* | * | ** | ** | ns | ns | * | ns | * | - | - | - | - | - |
| *A. uniaristata* | *** | *** | ns | *** | *** | ns | *** | *** | *** | - | - | - | - |
| *A. ventricosa* | ns | * | *** | ns | ns | *** | ns | ns | ns | *** | - | - | - |
| *H. vulgare distichon* | ns | ns | *** | ns | *** | *** | *** | * | *** | *** | *** | - | - |
| *T. dicoccum* | ns | * | *** | ** | *** | *** | *** | *** | *** | *** | *** | ns | - |
| *T. monococcum* | ns | ns | *** | ns | *** | *** | *** | * | *** | *** | *** | ns | * |

##### Subtype Ia

A few LM and M/N individuals (for example, HV11 and 16, H53, 64.x11, H327, H232) yielded a combination of oval A-Type granules and uniformly small, round B-Type granules (Fig. 4m). Our previous claims that this pattern is a recognizable feature of the domestic species of the tribe Triticeae (e.g., *Triticum* spp./*Hordeum* spp.) (Figs. 6,9) (Cristiani et al., 2016) are now further supported by a new morphometric analysis of both domestic and wild Triticeae species (Fig.9). Moreover, in the same individuals, A-Type granules could show cratered appearance (Fig. 4y). Similarly, in the EN individuals, lenticular and oval/sub-oval A-Type granules with equatorial groove and, an often visible, cratered surface are always associated with very small and uniformly shaped B-Type granules. Type A granules appear damaged in few EN individuals, which may be linked to enzymatic digestion (salivary amylase) although plant food processing could also result in starch damage based on experimental results (Ma et al. 2019; Zupancich et al. 2019)

**Fig. 4.**
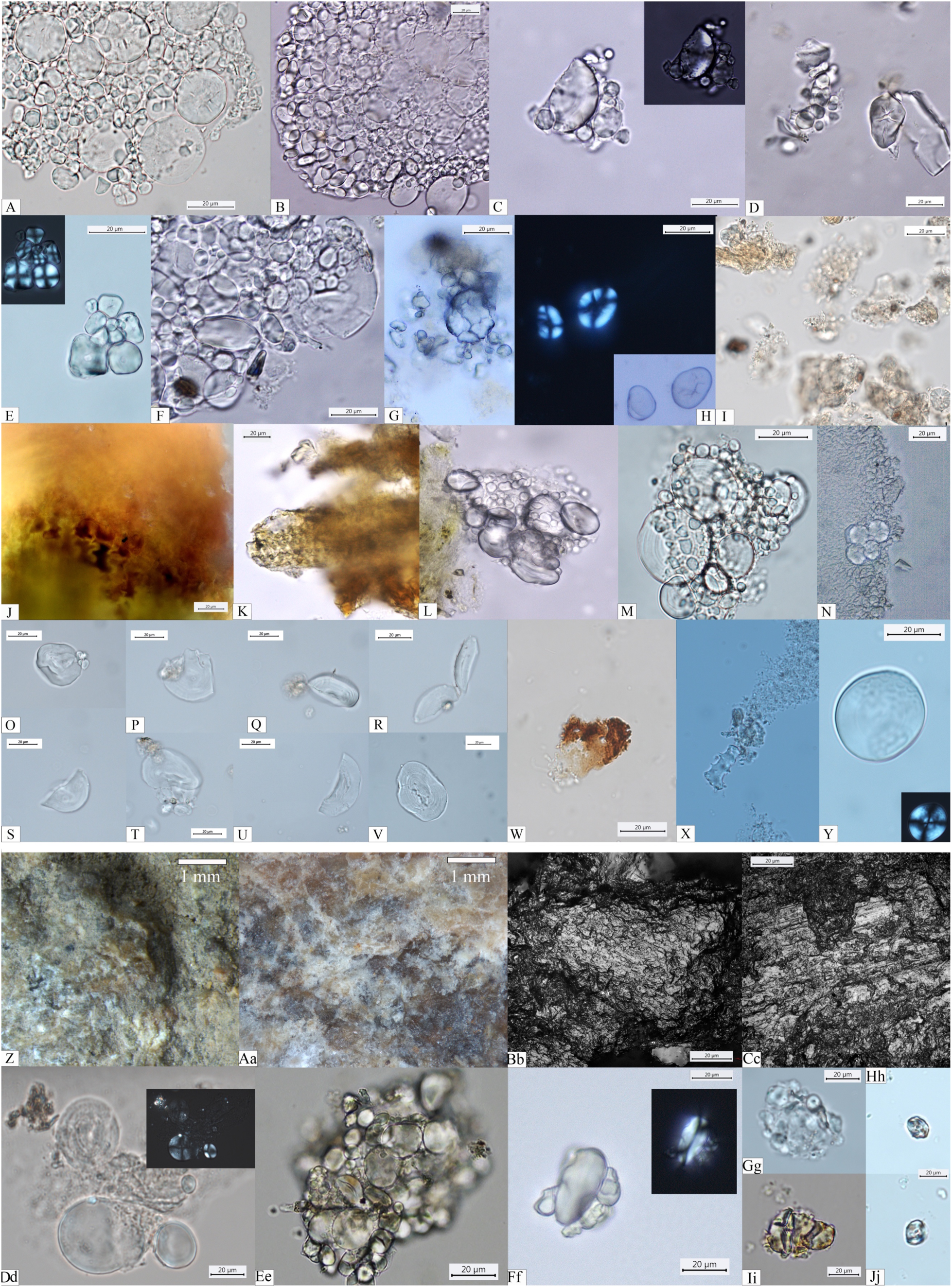
Starch granules from Mesolithic and Neolithic dental calculus (**a-y**) and functional evidence from ground stone tools (**z-Jj**). *Early Mesolithic:* (**a**) Type I (PAD11); (**b**) Type I (PAD9); (**c**) Type I (PAD11); (**d**) Type V (PAD12); (**e**) Type III (PAD11); (**f**) Type I (PAD15); *Late Mesolithic:* (**g**) Type II (VL82c); (**h**) Type IV (VL31); (**i**) Type VI (VL70); (**j, k**) Multicellular structures of long cells embedded in dental calculus (HV25/26, VL70); (**l**) Type I (HV11); *Mesolithic-Neolithic:* (**m**) Type I (HV16). *Neolithic*: (**n**) Type III (LV32a); (**o-v**) Damaged Type I granules (VEL-2D); (**w**) Multicellular dendritic structures (VEL-2D); (**x**) Single dendritic cell (Gîrlestj); (**Y**) Type I (VEL-2A). *Ground stone tools*: (**z**,**Aa**) yellowish organic residual film, covering the crystal grains across the utilized surface area on archaeological GSTs from Vlasac (GST no. INV80 and no.INV134).; (**Bb, Cc**) micro polish identified across the utilized surface of archaeological GSTs from Vlasac; (**Dd**) Type I (GST no. INV80); (**Ee**) Type I (GST no. INV146); (**Ff**) Type I (GST no. INV.71; (**Gg**) Type III (GST no. INV 28); (**Hh**) Type VI (GST no. INV 10); (**Ii**) Type VI (GST no. INV146; (**Jj**) Type VI (GST no. INV.67)

##### Subtype Ib

Significant unevenness in granule dimensions and shape was recorded in the EM and most of the LM individuals (e.g., PAD9 and VL45). Granules in this subtype are dimensionally variable and their shapes can range from round to oval (Fig. 4b). The well-known limitations in the inclusion, preservation, and recovery of plant debris in dental calculus (Radini et al., 2017) might be responsible for not recognizing this subtype previously.

#### Type II

Starch granules attributed to this type consist of large aggregates as well as clustered polyhedral/irregular granules (main axis ranging from 5 to 15 μm). They were retrieved from 10 individuals (2 EM, 11 LM, 2 M/N, and 3 EN) (Table 2) (Fig. 4g). The identification of archaeological specimens is based on published records (Lippi et al., 2015) and our experimental reference (*Avena barbata* L., *A. strigosa* Schreb., *A. fatua* L.) (Fig. 7). Granules of this morphotype were grouped in the tribe Aveneae/Poeae based on the fact that such large aggregates are found mostly in the genus *Avena* L. (oat), which is very common in the region.

#### Type III

Starch granules attributed to this type are characterized by a polyhedral to sub-polyhedral 3D morphology, a central hilum, and fine cracks. They were recovered in 16 individuals (5 for EM, 16 for LM, 2 for M/N, and 1 for EN) (Table 2) (Fig. 4e, Fig.5b and c). These features are consistent with starch granules of the tribe Paniceae of the grass family Poaceae and very well known in ancient starch research (Madella et al., 2016). In our sample, starch granules assigned to type III reach 21μm of maximum width, which falls within the size range found in several species of *Setaria* spp., *Panicum* spp., and *Echinochloa* spp. (Lucarini et al., 2016) (Fig. 7). Small granules characterized by a round to sub-polyhedral 3D morphology, and a central open hilum have been attributed to the tribe Andropogoneae and are here described under the general group of “millets” as is common practice (Madella et al., 2016).

**Fig. 5.**
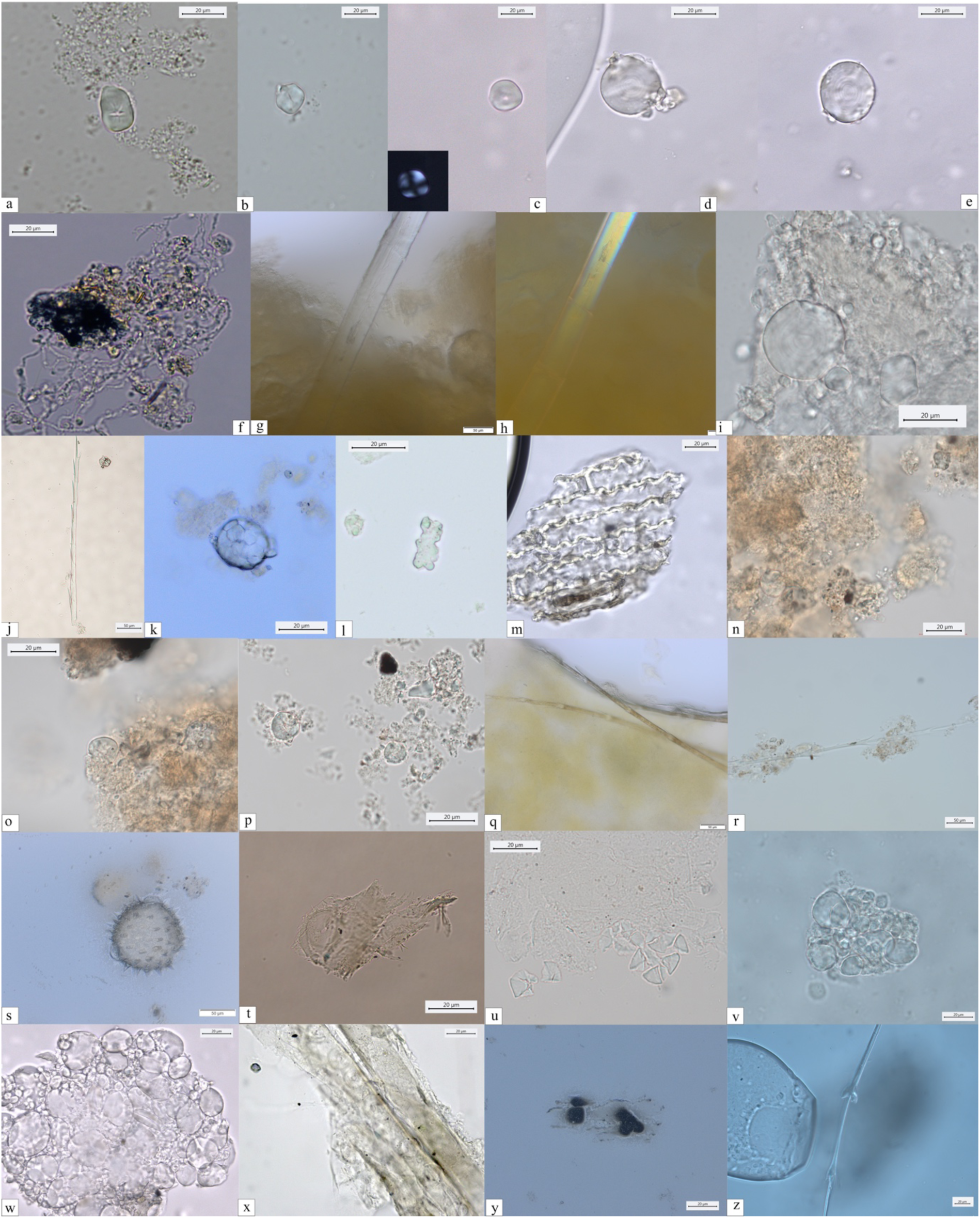
*Other dietary and non-dietary debris found in Mesolithic dental calculus from the Danube Gorges. Early Mesolithic:* (**A**) Type V (PAD11); (**B**,**C**) Type III (PAD12); (**D**) Type I (PAD12); (**E**) Type I (PAD12); (**F**) Smoke particle (LV50); (**G, H**) Plant fiber embedded in calculus (PAD16); (**I**) Type I (PAD9); (**J**) Feather barbule embedded in calculus (PAD9); *Late Mesolithic*: (**K**) Type II (PAD2); (**L**) Polylobate phytolith (US64 x.11); (**M**) Phytoliths (VL79); (**N-P**) Type VI (VL70,VL83); (**Q**) Feather barbules embedded in calculus (HV25/26); (**R**) Feather barbule embedded in calculus (VL80a); (**S**) Echinate pollen grain in calculus (VL83); (**T**) Wood particle (LV79a); (**U**) Type II (VL43); (**V**) Type I (HV11); *Mesolithic/Neolithic*: (**W**) Type I (HV16); (**X**) Wood particle (PAD4); (**Y**) Burnt phytoliths (LV28); (**Z**) Feather barbule (PAD4).

#### Type IV

Starch granules of this type are identified in 10 individuals (2 EM, 8 LM, 1 M/N, and 2 EN) and are recovered above 6 granules per specimen, with the exception of the only EN individual (Table 2). Diagnostic morphological characteristics for type IV granules are known in ancient starch research and include a reniform shape in 3D, a collapsed/sunken hilum forming a deep fissure along almost the entire granule, and a size ranging between 12 to 35 μm (Henry et al., 2011). Small cracks were observed departing from such hilum and were very evident under cross-polarized light. In most cases the extinction cross was very bright and showed several lateral arms diverging from the hilum in correspondence with the cracks (Fig. 4H). Moreover, lamellae were visible towards the outer part of the granules. All these features are very peculiar and diagnostic of starch granules included in the species of the plant family Fabaceae (Henry et al., 2011), which is mostly known for its several edible domesticated species of legumes (e.g. *Lens culinaris* Medikus, *Vicia faba* L., and *Pisum sativum* L.), but also has a number of wild edible such as vetches (*Vicia* spp.). While many edible species of the family Fabaceae grow in the Balkans (e.g. *Vicia sativa* L., *V. cracca* L., *V. hirsuta, V. ervilia, Lathyrus pratensis* L., *L. sylvestris* L.), an identification at species or genus was not possible due to overlaps in shape and size of starch granules at tribe level, which were observed in our modern reference collection (Fig. 7).

#### Type V

Few starch granules attributed to this type have been identified in 8 individuals (2 EM, 5LM, 1MN) (Fig. 4d, 5a and Table 2). Starch granules reach 23 μm in length and are mostly triangular with round corners and/or have an irregular oval shape (Fig. 5a). Overall, lamellae can rarely be visible. The granules show a linear fissure in the center and sometimes the hilum appears as a wide depression. Under polarized light, the hilum is mostly centric while the extinction cross has bent arms. This type was found to have a very close visual match with the starch found in acorns of oaks (*Quercus* spp.), a Fagaceae member and well known in ancient starch research (Liu et al., 2010) (Fig. 7i,j).

**Fig. 6.**
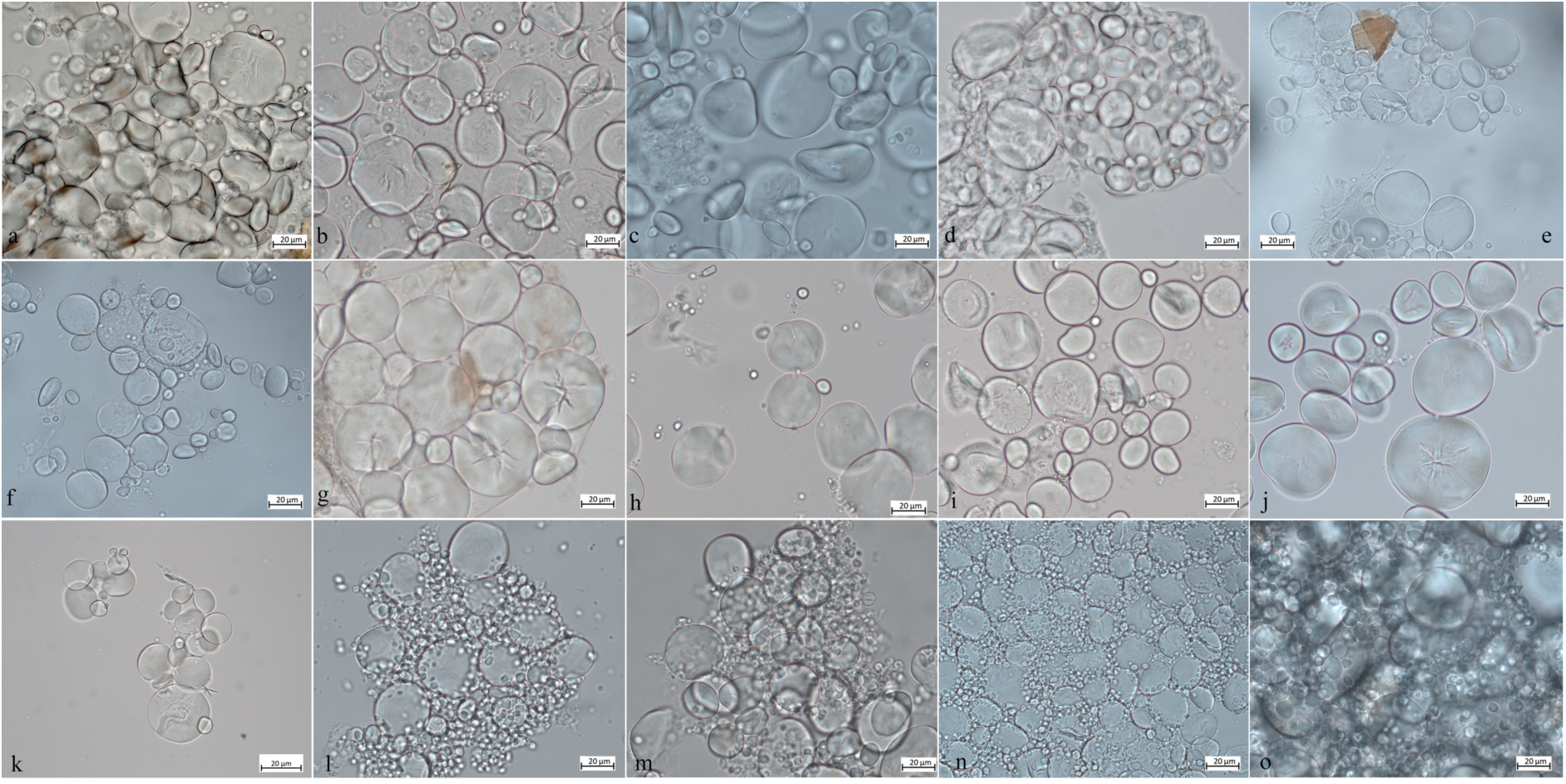
Starch granule morphological variability within the species of the genus Aegilops and domestic species of the Triticeae tribe. **A**) *Aegilops cylindrica*; **B**) *A. neglecta*; **C**) *A. speltoides tauschii*; **D**) *A. caudata*; **E**) *A. triuncialis*; **F**) *A. comosa*; **G**) *A. uniaristata*; **H**) *A. ventricosa*; **I**) *A. geniculata*; **J**) *A. crassa*; **K**) *A. peregrina;* **L**) *Triticum monococcum*; **M**) *Hordeum vulgare*; **N**) *T. dicoccum*; **O**) *T. aestivum*.

**Fig. 7.**
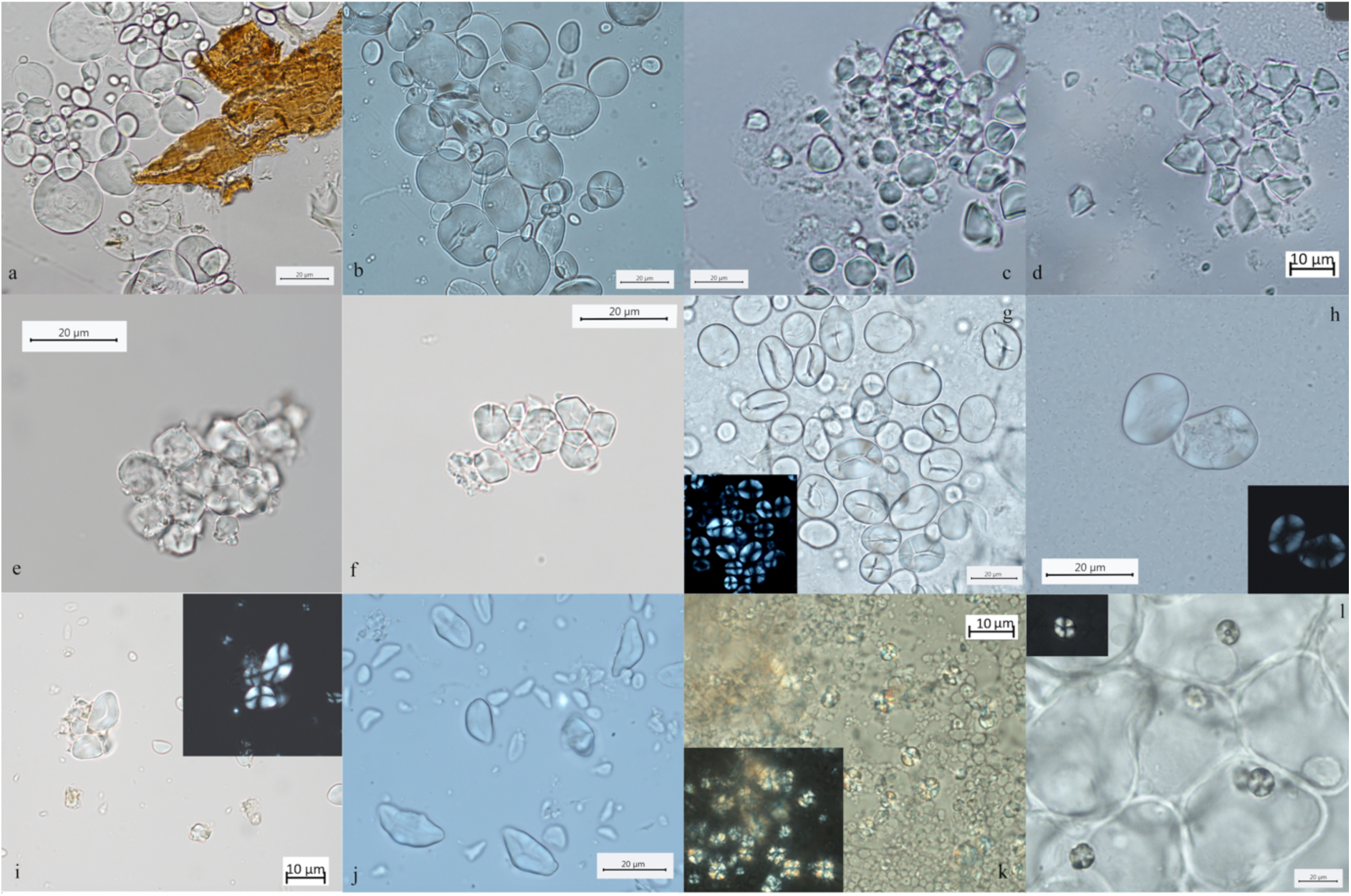
Modern reference used to interpret starch granules in archaeological dental calculus and GSTs. **A**) *Aegilops triuncialis*; **B**) *A. crassa*; **C**,**D**) *Avena strigosa*; **E**,**F**) *Setaria italica*; **G**) *Vicia cracca*; **H**) *V. sylvatica*; **I**) *Quercus pubescens*; **J**) *Q. robur*; **K**) *Q. colurna*; **L**) *Cornus mas*.

#### Type VI

Granules ascribed to this type have been identified in two individuals (1LM, 1M/N) (Table 2). They are characterized by a round 3D morphology and a central hilum, which appears as a wide depression, and no lamellae or facets (Fig. 4i, Fig. 5n-p). Zarrillo and Kooyman (2006) consider these morphological features diagnostic of some species of drupes and berries. In our sample, starch granules of this morphotype can reach 12 μm of maximum width, which is beyond species of berries and drupes in the Rosaceae family known in literature (Zarrillo and Kooyman 2006) and in our modern reference (e.g. *Prunus spinosa*). Based on our experimental record, we assign type VI to species of the family Cornaceae (e.g., *Cornus mas* L.) (Fig. 7), the remains of which are documented at Vlasac (Filipović et al. 2010).

In addition to starch granules, 42 phytoliths were retrieved in 24 individuals (EM=4; LM=13; MN=2; N=5). Mostly, short cells, commonly produced in leaves, stems and inflorescents, were identified (*14*) and attributed to Pooid grasses. Multicellular structures of long cells were identified in Mesolithic individuals (HV25/26, VL70, VL79, U222) (Fig. 4j,k; Fig. 5m). Of particular relevance is the recovery of multicellular phytoliths with dendritic appearance. It was observed at least in one case, still embedded in the dental calculus (Fig.4k). Single or multicellular dendritic structures were also identified in Neolithic individuals (VEL-2D and Gîrlestj) (Figs. 4w,x) (Fig. 4w). A single polylobate cell was found in one Late Mesolithic individual (US64 x.11). With the exception of dendritic structures, characterizing grass inflorescents, different non-dietary reasons may be suggested for the inclusion of phytoliths in dental calculus (i.e., accidental ingestion through food, breathing, flooring, etc.) (Norström et al., 2019).

### Ground stone tools

Diagnostic use-wear and residues are identified on 44 GSTs from the site of Vlasac. Analyzed tools included functional categories such as handstones (e.g., grinders and crushers) as well as passive bases (anvils) (Table 3). All of the tools are made of sandstone, characterized grains ranging in size between 0.2 and 1 mm densely distributed within the matrix. The combination of different functional modifications (i.e., flattened surfaces, pitted areas, rounding, etc.) on the single specimens suggests the long and complex life-histories of the artefacts, often used in different activities. Within the tools displaying diagnostic use wear, a total of 17 GSTs have surfaces bearing functional areas positively associated with plant food processing (Table 3 and Fig. 3). The analysis conducted at low magnification on these tools revealed macro-traces resulting in levelled surface crystal grains, sometimes covered by spots of yellowish organic film (sometimes striated) and white compacted powder (Fig. 4Z, Aa). At a high magnification, high and low micro topographies of the GST surfaces are affected by a smooth domed, and sometimes striated, micro polish (Fig. 4Bb, Cc). The aforementioned combination of use-wear and macro residues are commonly associated with GSTs used as handstones for crushing and grinding grass grains and/or fruits, such as hazelnuts and/or acorns in our experimental record (Cristiani and Zupancich, 2020) (Fig. 8).

**Fig. 8.**
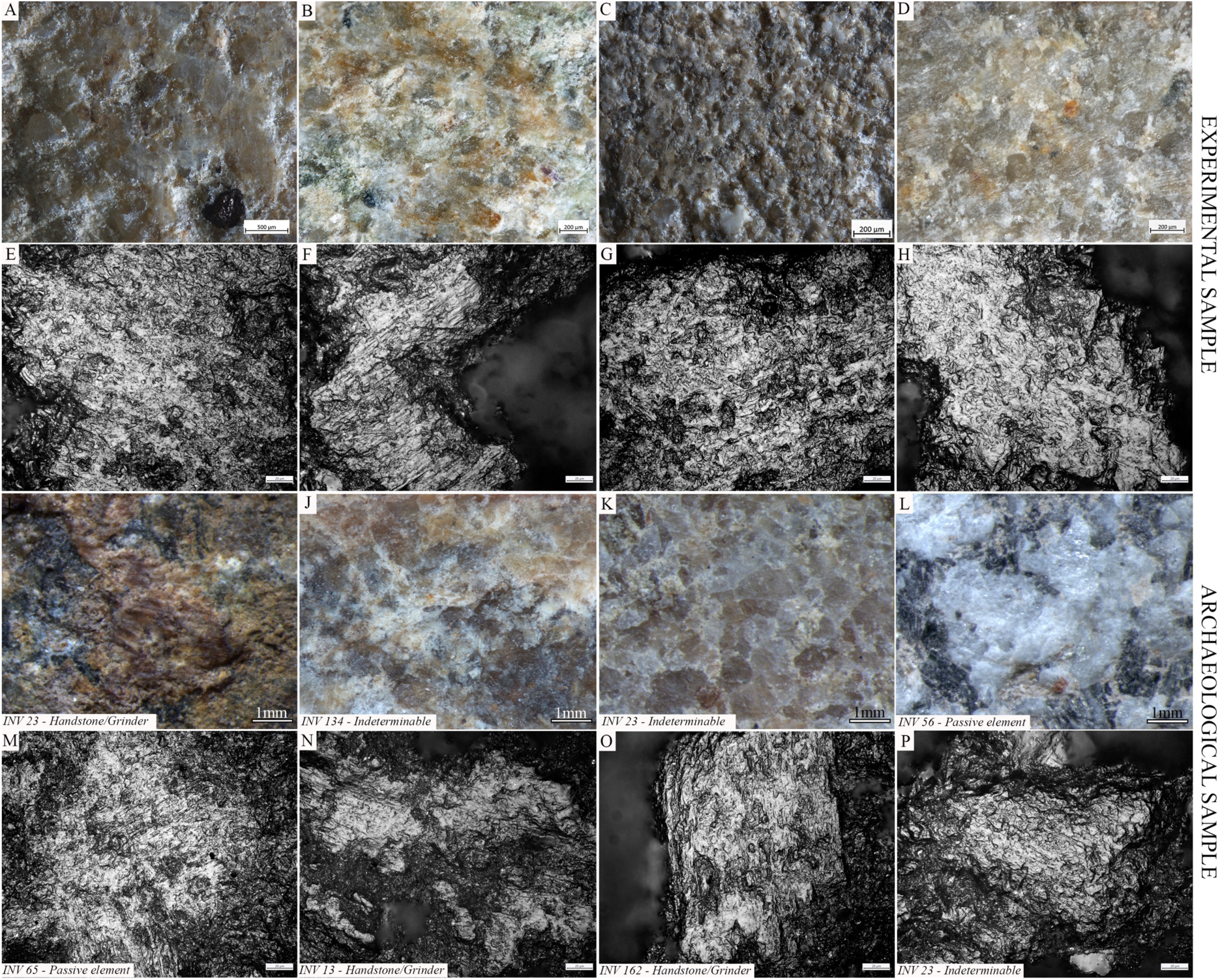
Experimental macro residues and micro polish associated with grass grains processing compared to macro residue and micro polish identified on archaeological GSTs from the site of Vlasac; **A-E**) yellowish organic film covering the crystal grains on experimental GSTs used to process oat (**A**), downy brome (**B**), wild grass grains, (**C**) and millet (**D**); smooth domed and flat micro polish developing over the high and low micro topographies associated with oat (*Avena barbata*) grinding; **F**) smooth flat and domed micro polish developing over the surface high and low micro topographies and characterized by narrow micro striations associated with grinding downy brome (*Bromus tectorum*); **G**) smooth flat micro polish developed over the high and low micro topographies characterized by sporadic narrow striations associated with grinding wild grass grains (*Aegilops ventricosa*); **H**) smooth domed polish developed over the high and low micro topographies associated with the grinding of foxtail millet (*Setaria italica*); **I-L**) spots of organic film, yellowish in color covering the crystal grains across the surface of archaeological GSTs; **M**,**P**) smooth domed micro polish identified on archaeological GSTs developing over the high and low surface micro topographies and associated with micro striations (**M**,**N**,**P**).

A total of 137 starch granules have been retrieved from the surfaces of the GSTs characterized by plant-related functional microscopic features. The optical and morphological properties of the starch granules support their attribution to morphotypes already documented in dental calculus from the Danube Gorges: type I assigned to seeds of the tribe Triticeae (*76*) (Fig.4 Dd, Ee, Ff); type IV, assigned to the Fabaceae family (*13*) (Fig.4 Ii); type III assigned to the tribe Paniceae of the grass family Poaceae (16) (Fig.4 Gg); and type VI, assigned to berries of the family Cornaceae (*32*) (Fig.4Hh;Jj).

In sum, several hundreds of starch granules and phytoliths of grass grains of the Triticeae tribe have been identified in the analyzed dental calculus of the Mesolithic population in the Danube Gorges. In addition, residues and use-wear identified on GSTs from Late Mesolithic Vlasac show the existence of a plant food processing technology during this period aimed at preparing a coarse-grained flour through a combination of pounding and grinding gestures (Table 3 and Figs. 3, 9). Grit particles, often retrieved in the analyzed dental deposit (Table 2), further confirm the use of sandstone GSTs in food processing. The conclusion about the consumption of partially processed grains is corroborated by the presence of starch granules still lodged in their amyloplast on Mesolithic GSTs and dental calculus, as suggested in our previous study (Cristiani et al., 2016). Interestingly, A-Type granules in EN dental calculus are generally retrieved singularly and exhibit a damage pattern observed when producing fine-grained flour experimentally only through prolonged bidirectional grinding (Dietrich et al., 2019). The pattern of bidirectional grinding is not documented on the examined Late Mesolithic GST from Vlasac, suggesting the existence of two different grain processing modalities typical of respective Mesolithic and Neolithic cultural traditions.

**Fig. 9.**
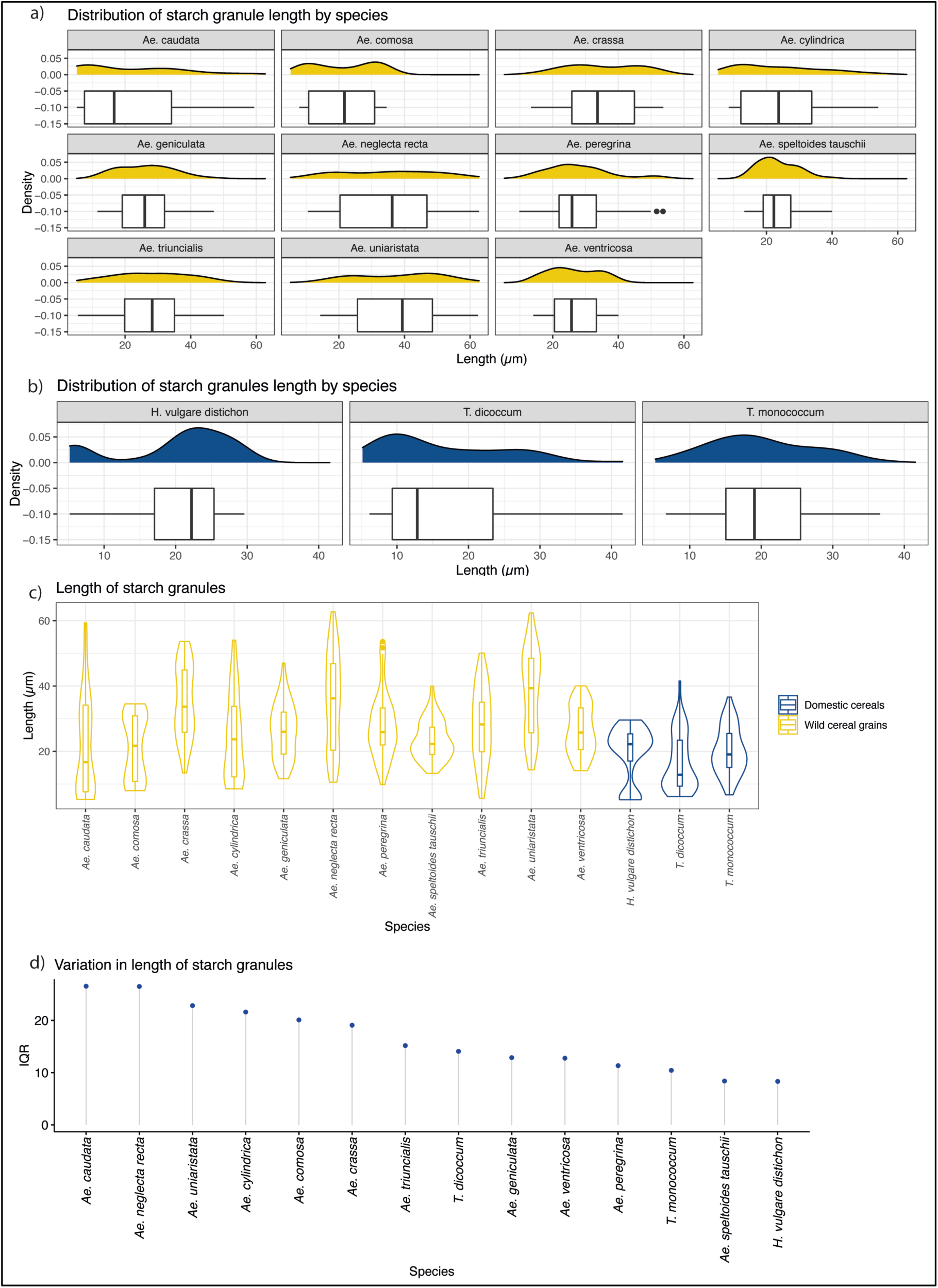
Starch granule length in modern wild and domestic cereal species. a) Distribution of starch granule length in wild species; b) distribution of domesticated species; c) Violin plot of comparing the length of starch granules in wild and domesticated species; d) IQR ranges of wild and domestic species. **Figure 9-source data 1**. Starch granule length in modern wild and domestic cereal species.

## Discussion

For some time now, there has been a recognition of the importance of plant foods in forager diets, based on both archaeological and ethnographic evidence (Clarke, 1978; Lee and DeVore, 2017). Research on mineralized dental plaque has significantly advanced our awareness about ancient pre-agrarian food choices in Europe, Asia and also in Africa, thanks to its potential to preserve plant micro-remains (Buckley et al., 2014; Cristiani et al., 2018, 2016; Cummings et al., 2018; Nava et al., 2021; Norström et al., 2019). However, in many archaeological cases dating to early prehistory, preservation or recovery biases render plant food evidence invisible. In exceptional cases, good preservation has allowed for the remains of plant macro-remains to be found, such as wild cereals at the Epipaleolithic site of Ohalo II in Israel, dating to 23 kya (Nadel et al., 2012; Piperno et al., 2004), or parenchyma remains at the Gravettian site of Dolní Věstonice in the Czech Republic (Pryor et al., 2013). On the other hand, micro-remains of oat caryopses have been detected on GST found at the Gravettian site of Paglicci cave in Italy (Lippi et al., 2015). In a seminal synthesis about plant foods in the European Mesolithic, Zvelebil (1994) reviews macro- and micro-botanical, palynological, artefactual (antler hoes, mattocks, GST), and human osteological (dental size and presence of caries) evidence for the consumption of nuts and fruits by European Holocene foragers, arguing for a form of niche constructing in temperate woodlands by means of deliberate forest clearance in order “to increase the productivity of nut and fruit trees and shrubs, wetland plants, and possibly native grasses” (Zvelebil, 1994). The emphasis is on the existence of some form of husbandry of wild plant species, which did not necessarily lead to domestication. Furthermore, between 200 and 450 indigenous European edible plants (grass seeds, nuts, fruits, roots, tubers, pulses) are found concentrated in wetland (coastal, lacustrine, and riparian) habitats (Clarke, 1978). Despite preservation and recovery problems, hundreds of Mesolithic sites across Europe have yielded the remains of hazelnuts, acorns, water-chestnuts, and other remains (Zvelebil, 1994).

In this paper, two complementary lines of evidence that we examined provide the first unambiguous and direct support for the consumption and processing of Poaceae grains among other types of edible plants by the Early Holocene foragers in the Danube Gorges area. The chronological framework of the analyzed sample suggests that this interest in and familiarity with various species of wild grasses of the Triticeae tribe (namely grass grains of the genus *Aegilops*) dates back to at least ~9500 cal BC. Macrobotanical remains belonging to this genus have not been recovered at Mesolithic and Early Neolithic sites in the central Balkans (Cristiani et al., 2016: 10301). However, such absence in the archaeobotanical record could be the result of a host of taphonomic and recovery problems and should not be used to exclude the use of this genus during the Mesolithic (contra Cristiani et al. 2016: 4). In fact, newly obtained evidence from the analysis of 61 individuals, which now also involves several EM individuals, lead us to suggest that *Aegilops* species were consumed in the region since the beginning of the Holocene. The mentioned difficulties associated with the recovery and preservation of botanical remains in local early prehistoric forager contexts along with some voids in the extant data regarding plant use by Mesolithic groups underlines the significance of our findings based on the application of relatively recent advances in dental calculus and GST analyses.

We have previously argued that three Late Mesolithic individuals from the site of Vlasac dated to the mid-seventh millennium cal BC (H53, 64.x11, and H232), as well as two presumed Early Neolithic individuals from Lepenski Vir (8 and 20) exhibit starches consistent with domesticated cereal species, such as *Triticum monococcum* L. (einkorn wheat), *Triticum dicoccum* L. (emmer wheat), and/or *Hordeum distichon* L. (barley) (Cristiani et al., 2016). This observation was based on the bimodal pattern of starch granules distribution, commonly attributed to domestic species and absent in most of the wild species of the genus *Aegilops* (Howard et al., 2011; Stoddard, 1999). This bimodal pattern is now further retrieved in two other individuals from two different sites (H327 from Vlasac and HV16 from Hajdučka Vodenica) dating to the Late Mesolithic and thus pre-dating the arrival of full-blown agriculture in the region. However, burial 20 from Lepenski Vir, previously published as dating to the Early Neolithic, has recently been directly AMS-dated to the Early Mesolithic (Table 1). This new chronological attribution does not correspond with our expectations that domesticated grains were introduced in the Danube Gorges only in the Late Mesolithic. At the face of the current evidence, we explain this inconsistency in our results by arguing that the admittedly small number of starch granules found in this archaeological specimen might have affected the visibility of the potential variation in A-Type *vs*. B-Type population. Moreover, a large variability of starch granule distributions among different species of the Triticeae tribe has been acknowledged in the literature (Henry and Piperno, 2008; Yang and Perry, 2013) and supported by our experimental reference (Fig. 9). Furthermore, fluctuations in environmental and growing conditions have also been recognised as relevant factors affecting starch granule size distribution (Stoddard, 1999; Stoddard and Sarker, 2000).

In addition to starch granules, other micro-remains, such as phytoliths and burnt debris, were recovered in the dental calculi of the analyzed Mesolithic population (Fig. 5L,M,Y). The paucity of archaeological phytoliths calls for caution when interpreting their dietary origin. Yet, the presence of few dendritic phytoliths in local forager dental calculus is likely related to plant consumption, as such micro-remains have experimentally been associated with mechanical destruction of husks and culms of Pooids by grinding (Portillo and Albert, 2014). In the investigated population, phytoliths provide also means of understanding the exposure to potential respiratory irritants generated during daily life activities (Radini et al., 2019, 2017; Scott and Damblon, 2010). The retrieval of phytoliths from herbaceous plants that appear burnt could suggest that plant materials, potentially harvested, were used in a diverse range of activities, including kindling and roofing. Phytoliths of such plants might have also been naturally present in the environment, water, and soil (Norström et al., 2019). Such burnt debris may have reached the mouth by accidental ingestion through food and/or breathing. Other micro-debris, such as nettle and wood fibers, might have also been linked to textile. Pathways to wood inclusion in calculus vary from the use of a toothpick to crafting activities (Radini et al., 2016) and the production and maintenance of weapons, such as arrowheads.

For the moment, it remains unclear to what extent plant foods, and species of the Poaceae family, contributed to the diet of the Danube Gorges foragers, and it is equally difficult to be more specific about the range of activities involved in their acquisition prior to processing and consumption – from sporadic collecting of wild grasses to forest clearances, or some form of indigenous system of plant management. Yet, it seems clear from our data that this specific subsistence practice was passed over generations in this regional context up to the first contacts of these foragers with Neolithic, farming groups in the second half of the seventh millennium BC. Previously, we suggested that some sort of exchange between Late Mesolithic foragers and first Neolithic groups in the southern Balkans might have allowed for the introduction of the domesticated species of Triticeae in the Danube Gorges area ~6500 cal BC (Cristiani et al., 2016). Now, our extended study seems to suggest that this introduction of domesticated cereals, i.e. more productive cereal strains from southwestern Asia, was preceded by several millennia of collecting and consumption of native grass grains. Adoption of previously unknown plant foods is often facilitated by those local foods that in shape and taste resemble the new arrivals (Livarda, 2011). Although not tested in this study, incipient management practices on certain wild plant taxa have been documented among non-agricultural societies for environmental and/or cultural reasons, for protecting or promoting the relative abundance of a species, or for reducing the energy involved in its harvesting (Fuller et al., 2011; Smith, 2011).

Another recent study of dental calculus from the Danube Gorges area has reached a conclusion that domesticated cereals started to be used in this region only with the start of the Neolithic (Jovanović et al., 2021). In the same study, the individuals dated to the Early Mesolithic exhibit a high occurrence of starch granules, which we have shown to be compatible with a variety of wild species of the Triticeae tribe (e.g., genus *Aegilops*). The evidence of Aegilops consumption during the Early Mesolithic was disregard as an “implausible pattern”. However, our results, based on the combination of a robust sample of analyzed individuals, GSTs, and experimental data are in opposition to the conclusions of the mentioned study.

Most of prehistoric forager groups might have had regular access to plant nutrients and some sort of dependency on specific plants (Fuller et al., 2011). Accordingly, in our analysis of dental calculus and GSTs, we have shown that besides wild grasses, local foragers consumed oat, legumes, minor millet species of the genus *Echinochloa* and/or *Setaria* known as “forgotten millets” (cfr Madella et al., 2016; Weber and Fuller, 2008), acorns, and Cornelian cherries (Fig. 4; Tables 1,2). Even if secure identification to species or genus level was not possible in every case (cfr. Soto et al., 2019), the morphological characteristics of starch granules systematically encountered in our archaeological samples, and the data available for the analysis of material culture, clearly show a contribution to the diet from these plant taxa.

Several starch granules of the tribe Aveneae have been identified in dental calculus, especially during the Late Mesolithic. Very abundant in temperate ecosystems, oat caryopses have been processed as foodstuff already during the Upper Palaeolithic at Paglicci cave, in the south of Italy (Lippi et al., 2015). They have also been documented in the dental calculus of the previously aforementioned Late Mesolithic individual from Vlakno Cave in the Eastern Adriatic region (Cristiani et al., 2018). Overall, starch granules of this tribe were remarkably well preserved in our dental calculus samples and large aggregates characterizing Aveneae were abundant. Interestingly, starches of this tribe recorded on archaeological GSTs, and associated with the production of flour, consist of abundant single sparse polyhedral granules (Lippi et al., 2015). In our case study, exceptionally well-preserved aggregates of Aveneae have been identified in dental calculus only. The concentration of semi-open aggregates (Figs. 4G, 5K,U) sustains the hypothesis of a coarse processing of the seeds during the Mesolithic, already suggested for other grass grain taxa.

Small starch granules from grasses of the Poaceae family have been attributed to the Andropogoneae and Paniceae tribes (Fig. 5B,C) (Madella et al., 2016). Our results confirm previous conclusions about the use of grasses of the tribe Paniceae by the Late Mesolithic foragers (Cristiani et al., 2016), while extending the consumption of this general types of millets to Early Mesolithic foragers too. However, no evidence of this food has been ascertained on the basis of stable isotope analysis (Borić et al., 2014). This pattern could mean that their consumption was not predominant in dietary practices of these Mesolithic foragers. We stress that a great variety of species of C4 plants belonging of the tribe Paniceae and the tribe Andropogoneae (e.g., species of plants of the genera *Setaria* Beauv. and *Echinochloa* P. Beauv.) grow well nearby water environments and slow-flowing waters, and it is very likely that a mixture of species from such genera might have contributed to the Mesolithic diet. Furthermore, they often grow in association with each other. The presence of Paniceae and the Andropogoneae tribes, combined with the evidence of several feather barbule fragments from aquatic birds, clearly point to a familiarity with resources from riverine environments other than fish.

Species of the family Fabaceae are well represented in the Early Mesolithic and Late Mesolithic individuals from the Danube Gorges area. While a secure identification to species or genus was not possible in our samples, we know that wild pulses (*Lens sp. Mill*.) and bitter vetch (*Vicia ervilia* (L.) Willd.) were used as foodstuffs for the Mesolithic inhabitants of Franchthi Cave (Asouti et al., 2018; Van Andel et al., 1987). Further evidence for a pre-Neolithic consumption of the species of the tribe Fabeae comes from Uzzo Cave, Italy, where wild legumes (*Lathyrus* sp. L., *Pisum* sp. L.) are well represented along with other arboreal fruits (*Arbutus unedo* L.), acorns (*Quercus* sp. L.), and wild grapes (*Vitis vinifera* subsp. *sylvestris* L.) (Costantini, 1989), as well as from the site of Barma Abeurador in southern France (Vaquer et al., 1986). In addition to the evidence from dental calculus, indirect evidence for the consumption of species of the family Fabaceae has also been retrieved from one GST (Fig. 4Ii).

Species of the Fagaceae family (cfr. *Quercus ilex* L., *Quercus* sp. L.) were also a significant food resource for the Danube Gorges foragers (Fig. 5A). Fragments of acorns were found at several Mesolithic sites across Europe (Holden et al., 1995; Kubiak-Martens, 1999) and in the investigated region (Mason, 1995).

Evidence for the consumption of Cornelian cherries (*C. mas* L.) has been retrieved in dental calculus belonging to Late Mesolithic and Mesolithic/Neolithic individuals from Vlasac (Fig. 5N-P) and also on eight GSTs from the same site. The results based on the analyses of GSTs not only corroborate the evidence for the use of these fruits derived from the calculus analysis, but also support the role of such plants in Mesolithic daily life. Cornelian cherries are the most recurrent macro-remains at Vlasac and are documented across Europe as a Mesolithic source of food and medicine (Divišová and šída, 2015; Filipović et al., 2010). Ethnographically, various fruits commonly referred to as “cherries” and/or “berries” are particularly well documented among Native American groups (Siegfried, 1994; Zarrillo and Kooyman, 2006). Throughout much of North America several species of the Cornaceae and Rosaceae plant families were processed using unmodified pounding stone tools to add to pemmican or to dry as cake. Interestingly, Woodland Cree people used to mix berries and cherries with fish eggs (Siegfried, 1994) and meat (Lowie, 1922).

We conclude that the long-lasting interactions with edible grains (but also wild pulses), documented in the Balkans since the end of the Pleistocene might have allowed enough time for specific eating habits, tastes, and “cultural valuation” (Hastorf and Foxhall, 2017) to develop. Such a shared knowledge about specific plant resources effectively predates the introduction of agriculture in Europe, and might have eventually eased the introduction of domesticated species starting from the second half of the seventh millennium BC. Our results call for more systematic and interdisciplinary research in order to reconstruct plant food traditions and cultural tastes before the introduction of agriculture.

## Materials and Methods

The examined collection of the teeth previously studied for strontium isotopes (Borić and Price, 2013) from five sites in the Danube Gorges area and one site from the region of Oltenia in Romania and additionally collected teeth from the sites of the Velesnica and the 2006–2009 excavations at Vlasac contained a total of 155 specimens. Of these, a total of 61 individuals had sufficiently preserved calculi for further analyses (Table 1): 13 individuals date to the Early Mesolithic (EM), 29 to the Late Mesolithic (LM), 9 to the Mesolithic-Neolithic transition phase (M/N), and 8 to the Early Neolithic (EN). In addition, two later period burials were also included in the analysis as a methodological comparison (Table 1). In order to corroborate data obtained through dental calculus analysis, we also analyzed 101 sandstone GSTs of the 131 implements found at the site of Vlasac during the 1970–71 excavations and chronologically attributed to the Late Mesolithic (Borić et al., 2014; Srejović and Letica, 1978). While GSTs are well documented during the Mesolithic-Neolithic transition at the site of Lepenski Vir (Antonović, 2007), earlier evidence for their use is available only from the site of Vlasac. This site yielded the richest Mesolithic assemblage of non-flaked stone tools recovered so far in the region (Antonović et al., 2006; Borić et al., 2014; Srejović and Letica, 1978). GSTs underwent a functional study aimed at verifying their function and potential involvement in plant food processing (Table 3 and Fig. 3).

### Dental calculus – sampling, extraction, decontamination and examination procedures

All the sampling was conducted under the stereoscope, as can be seen in Fig. 2 (for more details see below), and following strict protocols systematized by Sabil and Fellow Yates (Sabin and Fellow Yates, 2020) with some variation (disposable blades were changed after each sample extraction). Deposits of dental calculus were judiciously left on the teeth for future research. Whenever possible, sampled dental calculus was further subdivided for metagenomic analysis aimed at reconstructing aDNA of oral bacteria (Ottoni et al., 2021).

Decontamination and the extraction procedures for micro-debris were conducted in dedicated clean spaces not connected to modern botanical work and under strict environmental monitoring of the DANTE laboratory of Sapienza University of Rome (IT), the BioArch laboratory at University of York (UK), and the aDNA facility of the University of Vienna (Austria). In all of these facilities, strict contamination rules were followed. Cleaning is carried out daily and no food is allowed in order to prevent any type of modern contamination. Bench space surfaces were cleaned prior to the analysis of each sample, using soap and ethanol, followed by covering of the surfaces by aluminium foil, and using of clean starch-free nitrile gloves at all times. Calculus cleaning was carried out under the stereomicroscope, on a Petri dish previously washed and immersed in hot ultrapure water, with magnifications up to 100x. The removal of the mineralized soil adhering on the surface of the calculus was meticulously carried out using sterile tweezers to hold the sample and a fine acupuncture needle to gently scrape off the soil attached to the external layer of the mineralized plaque. The procedure was performed using drops of 0.05 M Hydrochloric (HCl) acid to dissolve the mineralized flecks of soil and ultrapure water to block the demineralization, as well as to wash and remove the contaminants. Once the calculus surface was cleaned, the contaminated soil was checked for possible cross contamination and the clean samples were washed in ultrapure water up to three times in order to remove any trace of sediment. The clean calculus was then dissolved in a solution of 0.5 M HCl and subsequently mounted on slides using a solution of 50:50 glycerol and ultrapure water. Furthermore, control samples from the clean working tables and dust-traps were collected and analyzed for comparative purposes in order to prevent any type of modern contamination in these laboratories – this is a practice routinely done in our laboratories, even at times where no archaeological analysis occur, to allow a better understanding of the flow of contaminations through seasons. Our results based on this procedure show that synthetic and plant fibers and hairs, fungal spores and hyphae, palm and conifer pollens, insects’ debris; maize starch granules were detected twice while unidentified small starch granules were very rare; phytoliths and starch granules belonging to species of the Triticeae tribe were never recovered (Fig. 10). We did not retrieve any debris morphologically similar to any of the remains in the environmental control samples. Furthermore, starch granules amounted to a neglectable fraction of the laboratory ‘dust’ – suggesting it is extremely unlikely that an event of contamination of starch granules would occur in the lab, where no other remains from dust, way more common, were not found.

**Fig. 10.**
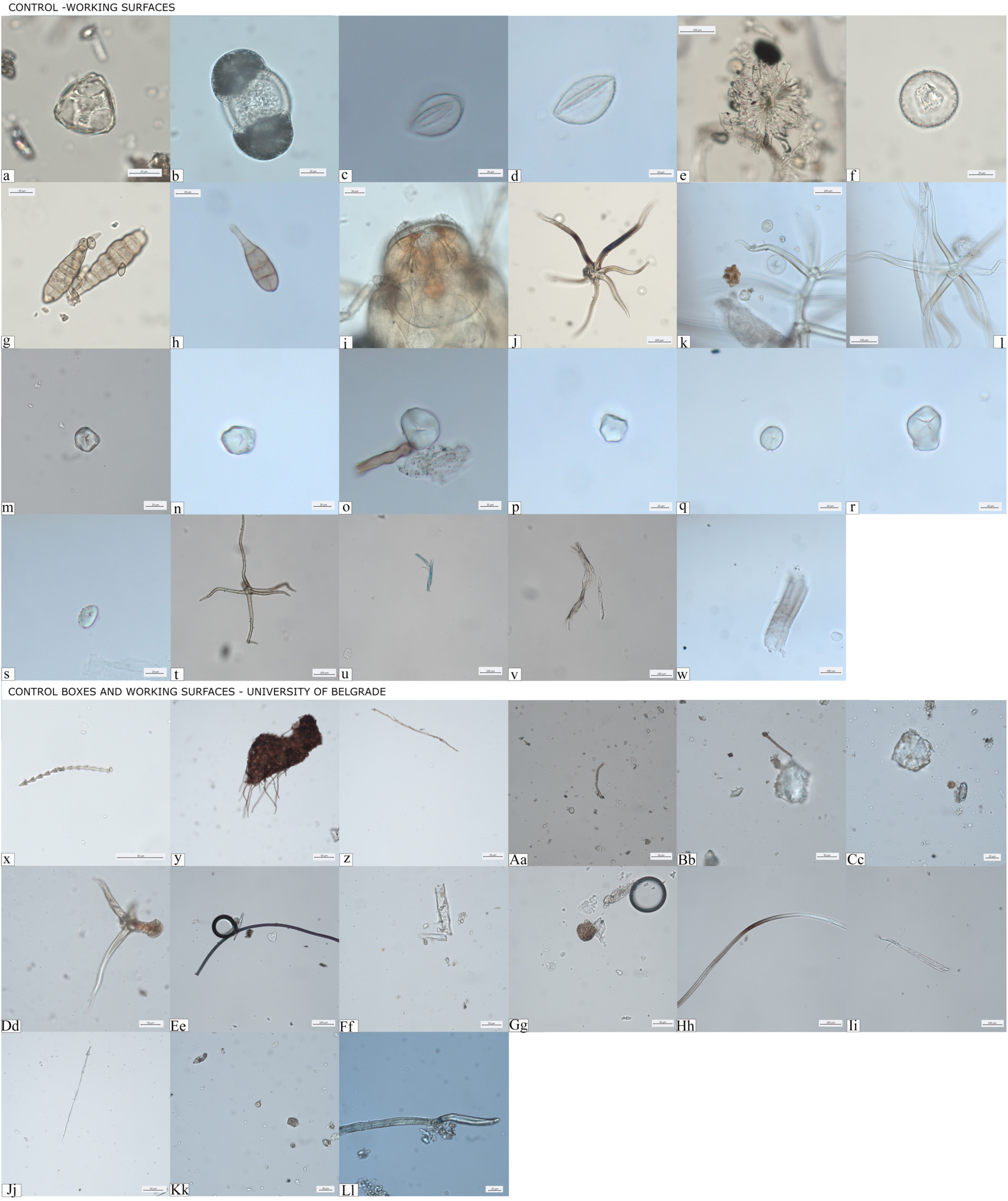
*Controls for contamination*. (**A-W**) Evidence of dust particles retrieved from clean working surfaces and dust traps located in different areas of the DANTE Laboratory at Sapienza University of Rome; (**X-Ll**) Dust recorded in storage boxes where groundstone tools were stored at the Faculty of Philosophy, University of Belgrade.

The examination of micro-debris embedded in the calculus matrix was performed at Sapienza University of Rome and at the University of York using a Zeiss Imager2 cross and an Olympus polarized microscope with magnifications ranging from 100X to 630X. A modern reference collection of 300 plants native to the Balkans, the Mediterranean region, and Europe was used as a comparison, along with published literature, for the identification of archaeological starch granules. The experimental reference collection also included species documented in the local archaeological record (Filipović et al., 2010; Marinova et al., 2013).

### Ground Stone Tools functional analysis

GSTs were sampled and analyzed at the Department of Archaeology of the University of Belgrade. Strict protocols were followed for controlling modern contamination during the residue sampling and functional analysis of GSTs: bench surfaces where the work was conducted were cleaned before the analysis of each tool using ethanol, hot water, and covered by aluminium foil; starch-free gloves were used while handling the GSTs; dust samples from the storage boxes and the working tables were collected and analyzed for comparative purposes; use-wear and residue analyses were performed on the surfaces not affected by severe post-depositional modifications and free from concretions. Furthermore, starch granules were considered reliable only when in combination with use traces associated to plant processing.

Functional study involved the analysis of use-wear traces on the GST surfaces at low magnification (0.75x–168x) using a Zeiss Discovery V20 stereomicroscope and at high magnification (200x– 500x) with a Zeiss AxioScope metallographic reflected light microscope(Cristiani and Zupancich, 2020; Dubreuil et al., 2015). At low magnification, GSTs were analyzed using a Zeiss Discovery V20 Stereomicroscope, which allowed us to assess the state of preservation of the materials and identify the residues still adhering to the surfaces. Appearance, morphological features, and spatial patterns of macro-residue distribution were considered (Langejans, 2010). Casts of the used areas were taken using a high-resolution polyvinylsiloxane (Provil Novo Light Fast Set), and later analyzed at high magnification (up to 500x) at the DANTE Laboratory at the Sapienza University of Rome. Micro-polish, abrasions, and micro striations across the tools’ surfaces were identified using a Zeiss AxioScope metallographic microscope, and described following relevant parameters available in literature (Adams et al., 2006; Dubreuil et al., 2015; Hamon and Plisson, 2009).

Micro-residues were sampled before surface casts. Ultrapure water was placed on the crevices of surface and left for one minute on the artefact in order to soften the residues, then pipetted out and stored in a sterile tube. Once in laboratory, the samples were centrifuged and the natant mounted on microscope slides using a 50% solution of purified water and glycerol. Slides were subsequently analyzed in transmitted light using Zeiss Imager2 microscope (630x) and cross-polarizing filters. For archaeological residues, appearance, morphological features, and spatial patterns of distribution were considered (Cristiani and Zupancich, 2020).

An experimental reference collection of used GSTs and starch granules housed at the DANTE laboratory was consulted along with relevant literature and scientific databases. High-resolution images of the identified use-wear and residue were taken at 630x using a Zeiss Axiocam 305 high definition color camera. Risk of modern contamination from the storage and sampling environment was minimized following a strict cleaning procedure before and during the sampling/analysis.

### *Testing morphological differences in experimental starch granules from* Aegilops, Hordeum, *and* Triticum *species*

Methodologically, we were able to characterize the morphological variability of starch granules in ancient plant species consumed in the investigated area, hence complementing our previous work and its implications (Cristiani et al., 2016). Differences in the starch granules assigned of the tribe Triticeae identified analyzed individuals were identified on the basis of (*1*) the specific morphology, dimensions, and appearance of type A and B granules; and (*2*) the proportion between A-Type and B granules. In particular, during the in Early Mesolithic (EM), A-Type granules preserved in calculi are very large and round in shape with deep lamellae visible only in the granules’ mesial part. Additionally, B-Type granules with different sizes and shape have been recorded in all of the EM individuals and most of Late Mesolithic (LM) individuals (Fig. 4). This feature is absent in the Early Neolithic (EN) individuals analyzed in this work, displaying only identical small round B-Type granules, while A-Type granules are large, round to oval/lenticular, with lamellae less pronounced in the mesial part of the grains and well visible craters on their surfaces (Fig. 4Y). We could not match these differences in our experimental record, which includes various *Aegilops* species as well as wild species within the genera *Hordeum, Elymus* L., *Agropyron* Gaertn., *Festuca* L. growing locally. The above-mentioned features have consistently been assigned to modern domestic Triticeae species (*Triticum* spp. and *Hordeum* spp.) (Cristiani et al., 2016; Piperno et al., 2004; Yang and Perry, 2013). Given the high variability recorded in the dimensions and distribution of starch granules within the modern, locally available, species of the genus *Aegilops* (Fig.6A-J), further statistical work was carried out in order to interpret starch granules assigned to the tribe Triticeae in Mesolithic/Neolithic transitional contexts.

Caryopses from 11 *Aegilops* species *(A. triuncialis, A. comosa, A. crassa, A. cylindrica, A. geniculata, A. neglecta, A. speltoides tauschii, A. peregrina; A. triuncialis, A. uniaristata, A. ventricosa)*, 1 *Hordeum* species *(Hordeum vulgare distichon)*, and 2 *Triticum* species (*Triticum monococcum*; *T. dicoccum*) grown in the central Balkans were collected. All plant material was grounded using pestle and mortar. Starch powder (0.5 mg) was resuspended in 100 µL of sterile distilled water and vortexed for 5 minutes. After that, the sample was observed by an optic transmitted light microscopy (Fig. 6). Fifty starch granules were randomly selected (for size and shape), counted, and their length measured. Minimum and maximum lengths, mean, and median values with relative standard deviations were reported for each species in Table 4. Length distribution and length variation of modern starches were reported in Fig. 9. Finally, length distribution of the starches from each species was compared with the other species to investigate the existence of significant differences. This statistical analysis was carried out through a Pairwise Wilcoxon test (Table 5). Results were considered significant for p-values <0.05 (<0.05; ^*^<0.01; ^***^<0.001) and not significant (n.s.) for measurements >0.05.

Triticeae starches are known to possess a bimodal distribution, made up of small (B-type) and large (A-type) granules (Fig. 6). In the analyzed *Triticum* and *Hordeum* genera, B-type grains are more abundant than A-type, except for *H. secalinum* Schreb., which does not exhibit the large granules. Differently, in *Aegilops* genus, the size distribution of starches is characterized by two different trends. The first one, evidenced in *A. caudata* L., *A. cylindrica* Host, *A. comosa* Sm., and *A. speltoides tauschii* Coss., appears very similar to that of *Triticum* and *Hordeum* samples, although B-type grains are less abundant than their counterparts in wheat and barley. On the other hand, the second cluster (*A. crassa* Boiss. ex Hohen., *A. geniculata* Roth, *A. neglecta* Req. ex Bertol., *A. speltoides tauschii* Tausch, *A. triuncialis* L., *A. uniaristata* Vis., and *A. ventricosa* Tausch) exhibits larger starches, determining a significant shift of the mean size towards intermediate lengths. In general, the present experimental analysis revealed that *Aegilops* sp. starch granules show a larger size distribution than *Hordeum* sp. and *Triticum* sp. (Figs. 6,9). This evidence is also supported by the Pairwise Wilcoxon test, which highlights that *Aegilops* spp. starch measurements are significantly different from those obtained for *Hordeum* sp. and *Triticum* sp. counts (Table 4).

## Acknowledgments

The authors wish to thank the late Živko Mikić (University of Belgrade), the late Borislav Jovanović (Serbian Academy of Sciences and Arts), Duško šljivar (National Museum in Belgrade), and Jelena Rankov (National Museum, Belgrade) for permissions to sample the osteological collections from the Danube Gorges.

## Author contributions

E.C. designed the study, collected and sampled all the data from prehistoric individuals, ground stone tools, and modern reference for starch analysis, analyzed all the data, contributed to the statistical analysis of the modern species of the Triticeae tribe, wrote the manuscript; A.R. analyzed part of the calculus samples and experimental reference, and wrote the manuscript; A.Z. analyzed functional data from ground stone tools, examined the experimental data, collected data and contributed to the statistical analysis of the modern species of the Triticeae tribe, wrote the manuscript; A. G. and A. D’A. contributed to the statistical analysis of experimental data, and contributed to the preparation of the manuscript, C. O. contributed to sampling and collected data; M.C. provided modern experimental reference for starch analysis, contributed to the experimental reference for ground stone tools; S. V. provided modern botanical reference and contributed to the preparation of the manuscript; D. A. and D. T. P. provided the samples and contributed to the preparation of the manuscript; D. B. contributed to the study design, collected all the data from prehistoric individuals and facilitated access to ground stone tools, provided chronological framework, wrote the manuscript. All the authors contributed to the preparation of the manuscript.

## Data and materials availability

All data needed to evaluate the conclusions in the paper are present in the paper and/or the Supplementary Materials. Additional data related to this paper may be requested from the authors.

